# Computation model predicts Rho GTPase function with the Plexin Transmembrane receptor GAP activity on Rap1b via dynamic allosteric changes

**DOI:** 10.1101/2025.03.13.643120

**Authors:** Nisha Bhattarai, Lindsay Morrison, Alexandre F. Gomes, Paul Savage, Amita R. Sahoo, Matthias Buck

## Abstract

Plexin-semaphorin signaling regulates key processes such as cell migration, neuronal development, angiogenesis, and immune responses. Plexins stand out because they can directly bind with both Rho- and Ras-family small GTPases through their intracellular domains when these GTPases are in their active, GTP-bound states. This binding occurs via intracellular regions which include a Rho-GTPase Binding Domain (RBD) and a GTPase Activating Protein (GAP) segment. Studies have shown that Rho and Ras GTPases play vital roles in plexin signaling and activation. However, the structural dynamics of plexins and GTPases and how these conformational changes affect interactions when plexin is bound with both Ras and Rho-GTPases or bound to only one specific GTPase has remained unclear.

In this study, we conducted molecular dynamics (MD) simulations on six distinct plexin-GTPase bound systems to investigate the differences in conformations and dynamics between Plexin-B1 and three GTPases: Rap1b, Rnd1, and Rac1. Our analysis revealed that dynamics with Rac1 are more altered, compared to Rnd1 depending on whether plexin’s GAP domain is bound or unbound to Rap1b. In addition, we further investigated alterations in network centralities and compared the network dynamics of the Plexin-GTPases complexes, focusing on the differences when Plexin is bound to both Ras (Rap1b) and Rho-GTPases (Rnd1/Rac1) versus when it is bound to only one GTPase. Our study revealed that Rnd1 exhibits stronger and more stable interactions with Plexin-B1 in the absence of Rap1b, while Rac1 shows fewer and less stable connections in comparison. These computational models have features that broadly agree with experimental results from hydrogen-deuterium exchange detected by mass spectrometry (HDX-MS). Such insights provide a better understanding of the molecular mechanisms underlying Plexin-GTPase interactions and the complexities of signaling mechanisms involving GTPases in general.

## 1. INTRODUCTION

Plexins are transmembrane proteins that serve as cell surface receptors for the axon guidance protein ligands, the semaphorins, in neuronal development but they are also involved in many other cell migration processes which are part of angiogenesis, immune,^2^ bone, and other developmental processes^1–4^. Several mutations have been identified in plexin receptors, leading to changes in protein function which then promote cancer^5–7^. Plexin proteins are characterized by a multidomain structure that includes an extracellular region, a single transmembrane helix, and intracellular domains^2,8^. The extracellular portion contains a ligand-binding Sema domain responsible for interacting with semaphorins^1^. What sets plexins apart is their ability to interact directly with small GTPases from both the Rho and Ras families segments of the intracellular region (ICR) – for plexin-B1 corresponding to residues 1511-2135^4^. The ICR consists of a juxta membrane region (JM), a Rho GTPase binding domain (RBD), and a GTPase-activating protein (GAP) domain^9–12^. The GAP domain shares structural similarities with RasGAPs, such as p120GAP, and contains key arginine residues analogous to the arginine fingers found in RasGAPs^13,14^. Semaphorin binds at the extracellular side for plexin signaling, resulting in receptor dimerization or at least a configurational change within the dimer, which is transmitted through the transmembrane region, promoting the activation of its cytoplasmic region, including the GAP domain^15^. Specifically, the GAP domain stimulates hydrolysis of GTP to GDP + phosphate for substrate Ras GTPases, such as Rap1b ^14,16^, but also stimulates the binding and regulation of plexin binding proteins, such as PDZ-RhoGEF and LARG.

The RBD (Rho GTPase binding domain) of plexin-B1 interacts with the Rho GTPases (Rnd1, Rac1, and RhoD) and is thought to be crucial for activating plexin’s GAP (GTPase-activating protein) activity in response to ligand binding and/or dimerization in cells^15,17–20^. In particular, Rac1 was initially believed to be an upstream activator of plexin, ^17^, but later it was proposed that plexin-B1 may be an effector of Rho GTPases. However, the binding of overexpressed Rnd1 to plexin is a more potent activator than Rac1^20^. Interestingly, while RhoD, Rac1, and Rnd1 both bind to plexins with similar affinity, only Rnd1 has been shown to stimulate plexins’ signaling activity in cells. ^20,21^

Several crystal structures have suggested that the RBD domain does not undergo substantial or global conformational changes upon binding with Rho-family GTPases^12,19,22^ however, how these changes differ in the presence of Ras-GTPases, particularly of Rap1b which has been identified as a plexin-GAP substrate, has remained elusive. Other protein-protein interaction systems showing dynamic allostery/extensive networking e.g. K-Ras:cRaf RBD^23,24^ have shown that possible changes can manifest in the conformational stability of protein subdomains and their internal dynamics, rather than in their static structures. Several studies have provided critical insights into the structural and dynamic aspects of Plexin-GTPase interaction^25–28^. For example, molecular dynamics (MD) simulations demonstrated that Plexin RBDs (Plexin-A1, -A2, -B1) exhibit isoform-specific interdomain dynamical networks when interacting with Rac1 and Rnd1^26^. A separate study also revealed the dynamic interplay between the JM domain and the activation switch loop^27^. The techniques used in this report, analysis of protein motions in molecular dynamics(MD) simulation trajectories^29–31^, and amide hydrogen-deuterium exchange, as detected by mass spectrometry,^32^ are amongst the few methods suitable to reveal such possible differences between Rnd1 and Rac1 binding to the RBD. Here, we study how such interactions and their dynamics are affected by the binding of the substrate Rap1b to plexins’ GAP domain.

To investigate this, we performed molecular dynamics (MD) simulations of Plexin-B1-ICR in different combinations of three GTPases (Rap1b, Rnd1, and Rac1). We ran MD simulations of six systems and analyzed their biophysical properties to uncover structural and dynamical differences between Plexin-B1 and the three GTPases in solution. We observed that Plexin-B1, in the absence of Rap1b substrate, has extensive cross-interface dynamic correlations (networks) between the RBD and Rnd1, whereas these are fewer in number for RBD bound Rac1. When Rap1b is bound, both RBD: Rnd1 and RBD: Rac1 networks are reduced in their extent and strength. Remarkably, these findings qualitatively match results from amide hydrogen-deuterium exchange as detected by mass spectrometry (HDX-MS). Overall, our data suggest that there is extensive dynamic coupling across the plexin-B1 intracellular domain but that this coupling can be altered by the binding of partner proteins in a protein-specific manner, which shows some correspondence with the different functional role of the Rho GTPases in the plexin signaling mechanism.

## 2. RESULTS

Two known plexin-GTPase interfaces were analyzed by all-atom molecular dynamics simulations of various complexes. **Figure 1a** shows the initial simulation system for the Plexin-B1_Rnd1_Rap1 (B1_Rnd1_Rap1) complex. **Figures 1b** and **1c** highlight the binding region between RBD and Rho GTPase and GAP and Rap1b, respectively. We prepared similar systems with Plexin-B1_Rac1_Rap1b, Plexin-B1_Rap1 only, Plexin-B1_Rnd1_only, Plexin-B1_Rac1_only, and Plexin-B1 only.

**Figure 1:**
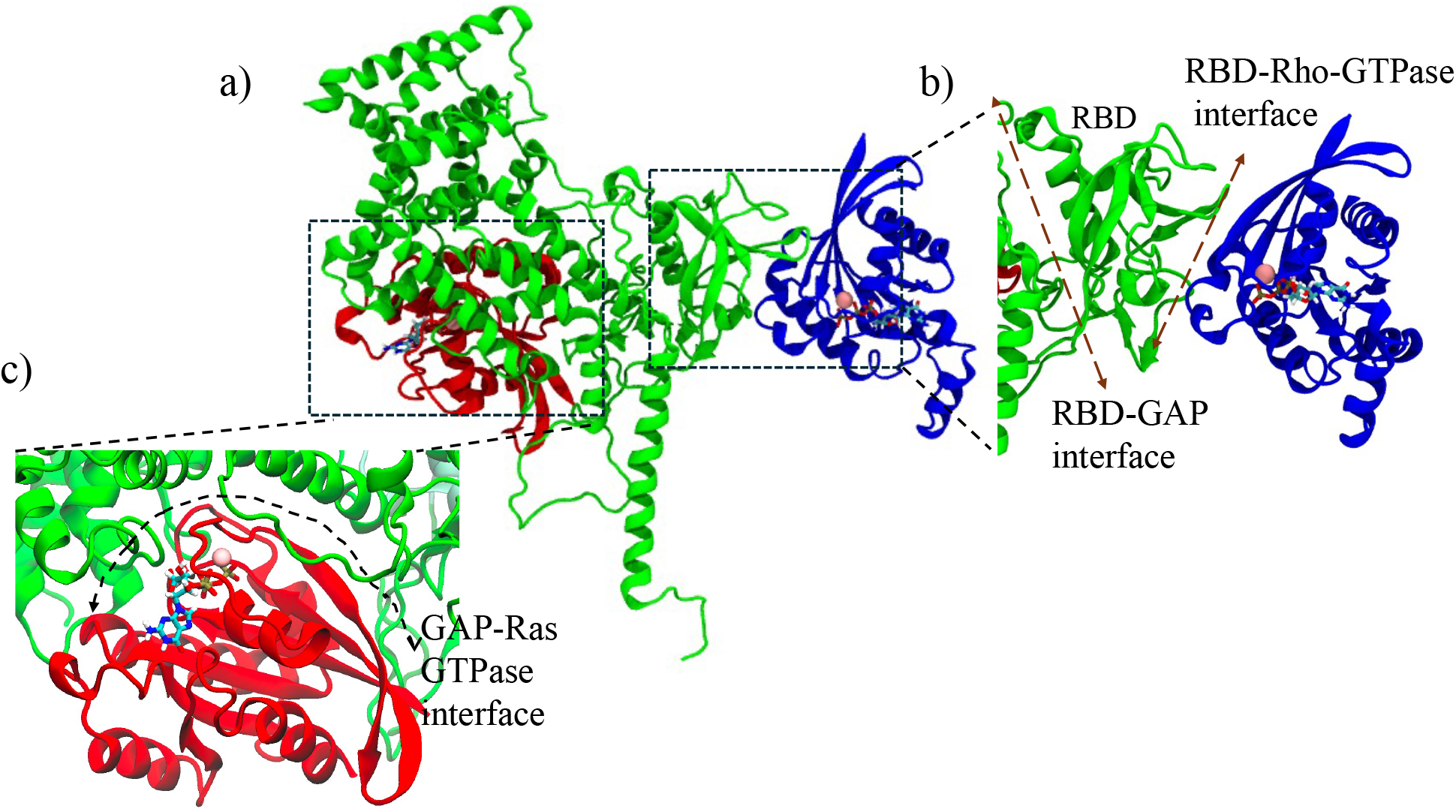
a) Initial system showing Plexin-B1-ICR in complex with Rho-GTPase (Rnd1) and Ras-GTPase (Rap1b), where Plexin-B1 is colored green, Rnd1 in blue, and Rap1b in red. GTP molecules are shown in the sticks, and Mg^2+^ ions are represented as spheres. Zoom in on the b) the binding region between RBD (dark green) and Rnd1 (blue), and c) the binding region between the GAP domain (green) and Rap1b(red).

### 2.1 Stability of all complexes

To evaluate the equilibration of each system during the simulation, we analyzed the root mean square deviation (RMSD) for all systems, examining all three replicas. The RMSD was calculated for the backbone atoms over a 200 ns trajectory for each replica, totaling 600 ns of simulation time. We separately analyzed and plotted the RMSD for Rap1, Rac1, Rnd1, and Plexin_B1 across all complexes. We then averaged the RMSD values from all replicas and plotted these averages with standard error as a function of simulation time, as shown in **Supplemental Figure 1**. For the Rho GTPases Rac1 and Rnd1, RMSD values stabilized at approximately 0.12 nm after 50 ns for all complexes (Rnd1_only, Rnd1_Rap1, Rac1_only, Rac1_Rap1). In contrast, for Plexin_B1, RMSD values stabilized around 0.4 nm for all complexes, except for the B1_Rac1_only system, where it reached approximately 0.5 nm. This suggests that Plexin_B1 might be slightly more flexible when interacting with Rac1 only in the absence of Rap1.

To analyze the conformational sampling of each protein complex, we used principal component analysis (PCA) on the covariance matrix derived from the carbon alpha atomic fluctuations over time. Specifically, the method projects the deviation of the backbone atoms into a subspace defined by the first two principal components (PC1 and PC2), representing the most extensive motions, aiding us to visualize how the proteins explore conformational space. We only considered the first two PCs (PC1 and PC2) as they represented more than 50% of the conformations. **Figure 2** illustrates the PC1 and PC2 projections for each protein complex across all three replicas, with each replica in a different color. The plots reveal that conformations from different replicas often overlap, indicating a robust sampling across different runs. Among the protein complexes, the B1_Rac1 complex explored the most conformational space, suggesting greater fluctuations. This was followed by the B1_Rnd1_Rap1 complex and the B1_Rac1_Rap1 complex, which also displayed considerable conformational variability. Interestingly, the B1_Rnd1 complex explored less conformational space compared to the B1_Rac1 complex. The B1_only sample, i.e. the uncomplexed protein, exhibited the least conformational variation, with the B1_Rap1 complex showing slightly more variability but still relatively constrained compared to the others.

**Figure 2:**
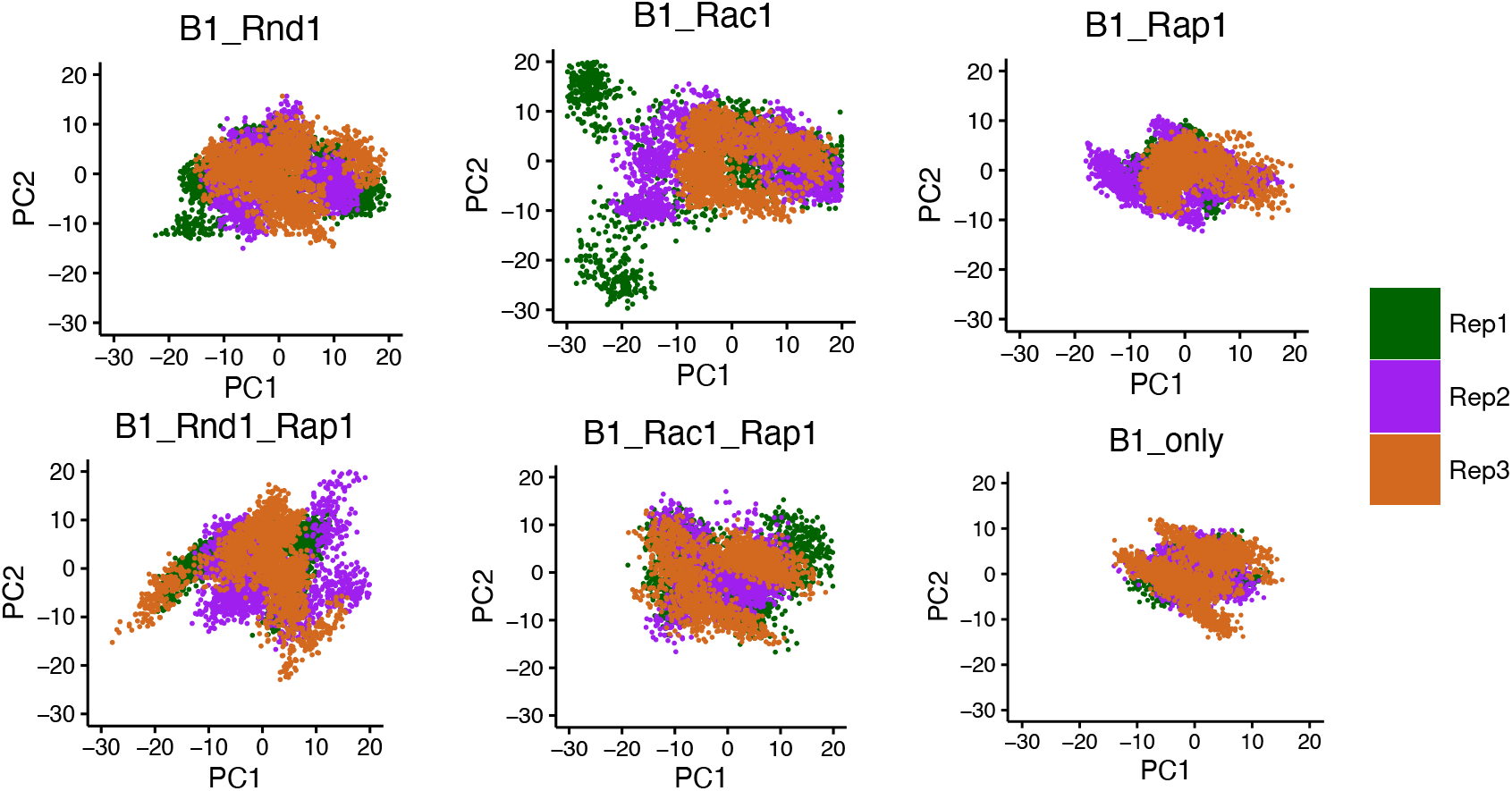
Projection of the Cα atoms of all six complexes on the essential subspace, defined by the first two eigenvectors (PCs) of the covariance matrix of the protein. The colors indicate the three different replicas (200 ns each), where Replica-1, -2, and -3 are colored in green, purple, and chocolate, respectively.

To gain further insight into the fluctuations within the complexes, we compared the RMSD plots with root mean square fluctuations (RMSF) and local frustration analysis (**Figure 3** and **Supplemental Figure 2, respectively**). We observed that the B1_Rac1_only complex exhibited greater flexibility overall, especially in the loop region spanning residues 1853 to 1913, which connects the RBD and GAP domains of Plexin-B1. This finding aligns well with the frustration analysis, where we noted that these regions were minimally frustrated when binding to Rap1b in the B1_Rac1_Rap1 complex, as compared to the B1_Rac1_only complex. For the Ras GTPase Rap1b, RMSD values were consistently stabilized at around 0.11 nm across all systems.

**Figure 3:**
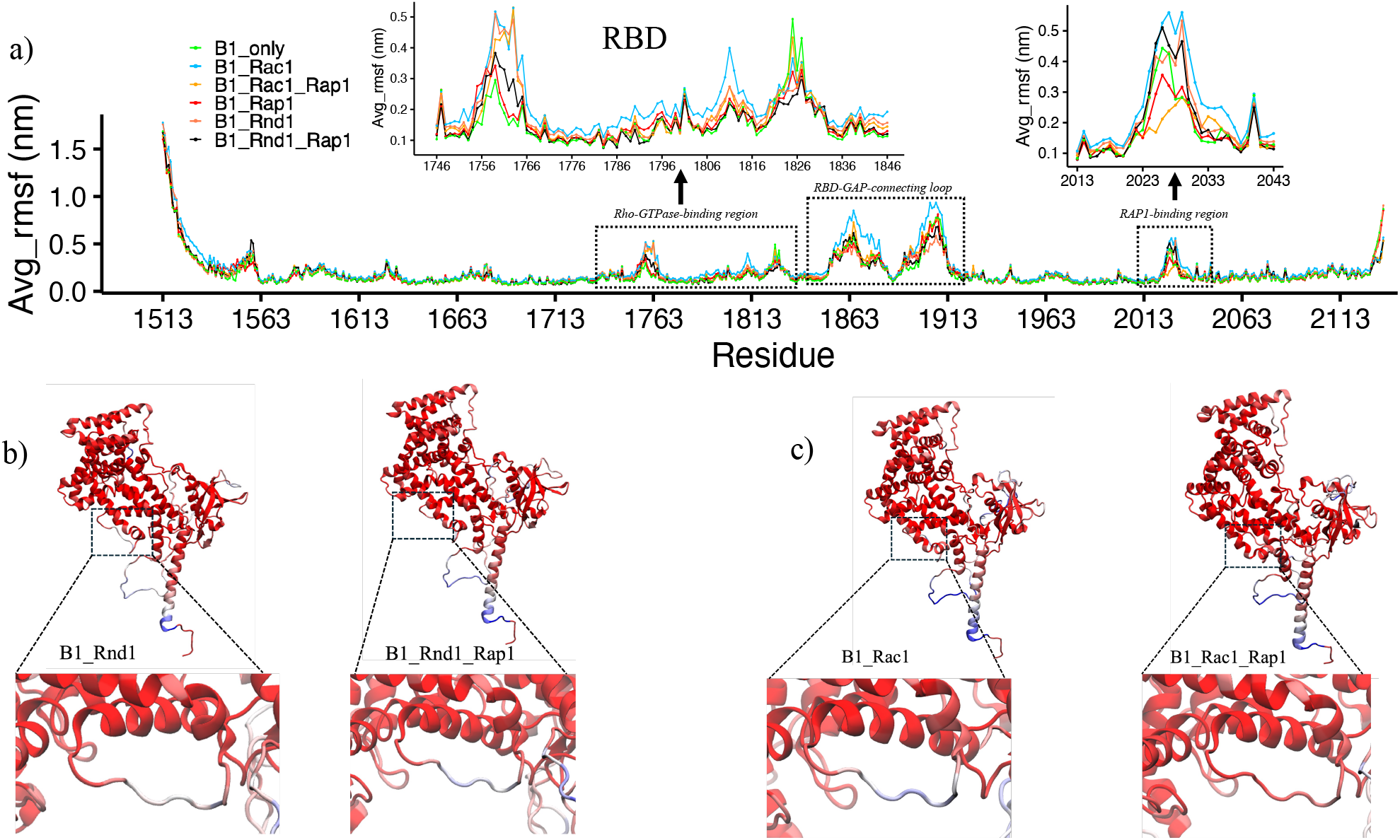
a) Root mean square fluctuations for Plexin-B1 for all six complexes (B1_only: cyan, B1_Rac1: green, B1_Rac1_Rap1:yellow, B1_Rap1:red, B1_Rnd1:coral, B1_Rnd1_Rap1:black). The first dotted rectangle represents the RBD region of Plexin-B1(inset: zoom in of RBD). The second dotted highlights the RBD-GAP domain connecting loop and the third dotted region shows the Rap1b binding section(inset: zoom in). b) and c) RMSF values color-coded on protein structure where the Rap1b binding region is highlighted.

### 2.2 Difference in Plexin-B1 Dynamics

We next focused on Plexin-B1 as a whole and studied how Rho/Ras GTPases binding changes internal Plexin-B1 dynamics, by –as above-again analyzing the same simulations of the six different systems (see Methods) simulated in solution with three replicas each of 200 ns. We mainly focused on the Rho-GTPase binding domain (RBD) and Ras-binding region (GAP-region) of Plexin-B1. In **Figure 3a**, we assess the flexibility of each residue in Plexin-B1 by plotting the average RMSF values of the three replicas. Plexin-B1 itself, without GTPases was the least flexible, as already seen in PCA (**Figure 2**). We observe significant changes in the Rho GTPases binding domain: RBD between the complexes. The RBD in B1_Rac1 was the most flexible, followed by B1_Rac1_Rap1 and B1_Rnd1 (inset: 1742:1846). Interestingly, with Rap1b, the RBD in B1_Rnd1 fluctuates less. Another region of interest, RBD-GAP connecting loop residues: 1853-1913, was most flexible in the B1_Rac1_only complex; however, upon binding with Rap1b, it is less flexible. On comparing with Rnd1 complexes, we observe that Rap1b binding affects Rac1 more than Rnd1, as these flexibilities were significantly different with Rac1 complexes. In addition, from local frustration analysis focused on the RBD-GAP domain connector, we observe similar findings: B1_Rac1_Rap1 was minimally frustrated and was significantly different than the B1_Rac1_only system. However, for the Rnd1 system’s local frustrations, there were not many differences **(Supplemental Figure 2)**.

Moving on to one of the Rap1b binding regions (inset: 2013-2043) of Plexin-B1, the results are different. This region is highly flexible when no Rap1b is bound (green, blue, red-orange) but experiences decreased fluctuations when interacting with the substrate GTPase, except, remarkably, when Rnd1 is bound at the RBD segment (black-) it has an extent of fluctuations similar to this GAP region without Rap1b. The B1_Rac1_Rap1 system was more stable than B1_Rnd1_Rap1, which is also highlighted in **Figures 3b** and **3c**. These differences were more subtle with the presence of Rac1 than Rnd1. We also note here that we did not observe significant differences in conserved arginine residues at the plexin GAP domain (R1677 and R1678), catalytically essential for Ras GAPs as mutating these residues abolishes plexin activity both in vitro and in vivo ^33^.

Although we did not observe any differences in Rap1b between B1_Rac1_Rap1 and B1_Rnd1_Rap1 in terms of network closeness between Rap1b and Plexin-B1, the difference was observed in Plexin-B1 flexibility with and without Rnd1 or Rac1. It is important to note that we also compared closeness centrality for Plexin-B1 overall and RBD region by itself **(Supplemental Figure 6)** but did not observe noticeable changes with Rho-GTPases with/without Rap1b. However, the residue flexibility in Plexin-B1 for B1_Rac1_Rap1 is very similar to B1_Rap1 and the fluctuations of residues were higher with Rnd1. Overall, these results suggest that Plexin-B1 dynamics are affected by the type of Rho-GTPase it interacts with and whether Rap1b is present.

### 2.3 Rho GTPase dynamics: Rnd1 and Rac1 dynamics with Plexin-B1 in presence and absence of Rap1b

Next, we examined the fluctuations of each residue within the Rho GTPase protein structures by calculating the root mean square fluctuations (RMSF) for all systems and replicas. We then averaged these RMSF values for each residue and plotted them to visualize the fluctuations.

**Figures 4a** and **4b** present line plots of RMSF for Rnd1 and Rac1, respectively. **Figure 4b** shows that RMSF values for Rac1 systems did not show significant differences except for slight changes in regions 31 to 40, which correspond to the switch region for the GTPase. For Rnd1, there were noticeable variations as well, specifically, RMSF values for Rnd1 were slightly higher in certain regions (residues 90-110) when plexin is also bound to Rap1b.To further illustrate these differences, we color-coded the average RMSF values onto the protein structures. **Figures 4c** and **4d** display the RMSF values for the B1_Rnd1 and B1_Rnd1_Rap1 complexes, while **Figures 4e** and **4f** show the RMSF values for the B1_Rac1 and B1_Rac1_Rap1 complexes. The observed minor differences are highlighted by a dotted circle, in the region spanning amino acids 90 to 110, which is located slightly away from the RBD binding region, for the Rnd1 complexes, and regions 31 to 40 for the Rac1 complexes, corresponding to the switch region of GTPases. In addition to RMSF, we also calculate interaction energy between Plexin-B1 and Rho-GTPases **(Supplemental Figure 3)** to check the stability of these complexes. The interaction energy for B1_Rnd1 systems was less than the B1_Rnd1_Rap1 system, but it was higher in the B1_Rac1 system compared to B1_Rac1_Rap1 system. These findings indicate that the effect of Rap1b is more pronounced for Rnd1 than Rac1 GTPase when interacting with Plexin-B1.

**Figure 4:**
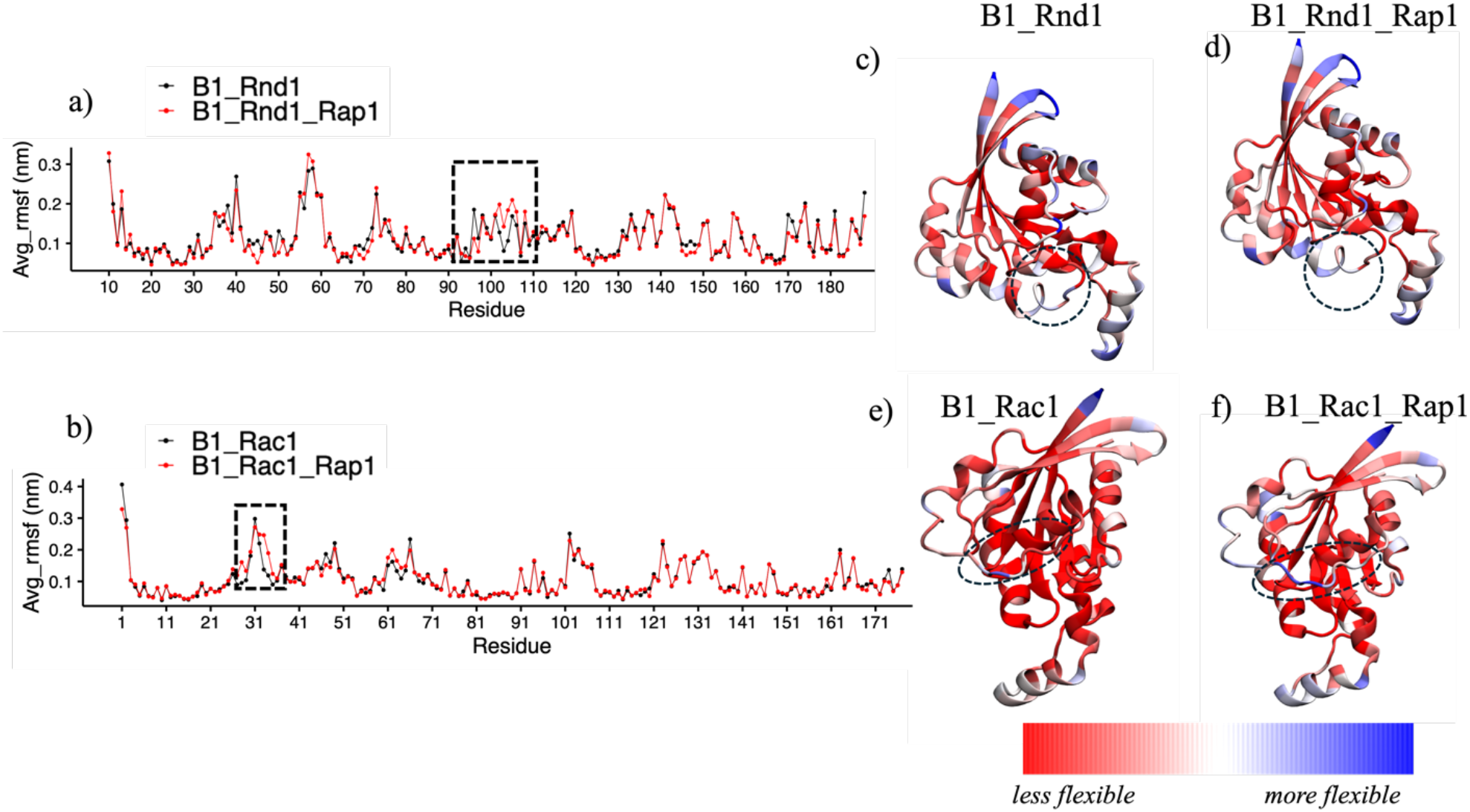
Root Mean Square Fluctuations (RMSF) Analysis of the GTPases a) RMSF data for Rnd1: B1_Rnd1 is depicted in black, while B1_Rnd1_Rap1 is shown in red. b) RMSF data for Rac1: B1_Rac1 is shown in black, and B1_Rac1_Rap1 is shown in red. Structural representations color-coded by average RMSF values (red-white-blue): Structure of Rnd1 c) B1_Rnd1 complex. d) B1_Rnd1_Rap1 complex. e) Structure of Rac1 e) B1_Rac1 complex f) B1_Rac1_Rap1 complex. The dotted circle indicates residues 90-110 in Rnd1 proteins and regions 31 to 40 on Rac1 proteins, where the minor differences between the two complexes were observed.

#### Network Analysis for Rnd1 and Rac1 dynamics

After observing these fluctuation changes in B1_Rnd and B1_Rac1 in the presence and absence of Rap1b, the question arises as to how Rac1/Rnd1 communicate with the plexin-RBD they are bound to and how these communications are different with/without Rap1b bound to plexins GAP domain. To explore this communication, we conducted a network analysis which will allow us to understand the topology and dynamics of residue interactions. For the network analysis, we first performed cluster analysis on all three simulation trajectories (600 ns total, combining three replicas). We extracted the conformation of the major cluster for each complex and used these conformations as input for network analysis via the NAPS webserver^34^.

Figure 5. illustrates the network connections between Plexin-B1 and Rho GTPases (Rnd1 and Rac1), as well as between Plexin-B1, Rho GTPases, and Rap1b. We observed a significantly higher number of connections in the B1_Rnd1 complex (∼15 connections) compared to the B1_Rnd1_Rap1 complex(∼7 connections, **Figure 5a**). In contrast, for the Rac1 systems, there were not many changes in number of connections in the B1_Rac1 complex (∼14 connections) compared to the B1_Rac1_Rap1 (∼15 connections) complex (**Figure 5f**). To gain further insights into which amino acids are involved in these interactions, we computed the contacts formed between each GTPase and Plexin-B1 across all complexes and visualized them as a contact map **(Supplemental Figure 4)**. This analysis was conducted using the Mapiya web server^35^ on major cluster conformations and plotted in R. For Rnd1, we observed a higher number of contacts with Plexin-B1 when Rap1b was unbound, which aligns with the increased network connections seen in **Figure 5**. However, for Rac1, this difference was not significant.

To validate these results, we performed additional network analysis, i.e., a dynamical network analysis using VMD. This analysis included all frames from the three replicas and focused on the carbon alpha atoms of the proteins. The dynamical network analysis corroborated the findings from the NAPS web server, showing that the B1_Rnd1 complex had more connections than the B1_Rnd1_Rap1 complex. Interestingly, for the B1_Rac1 system, no connections were detected, and only one connection was found in the B1_Rac1_Rap1 complex **(Supplemental Figure 5)**. After exploring overall protein-protein network connections, we dig deeper to characterize at the individual residue level how these protein complexes are communicating, and what amino acids play critical roles in these communications. We focused on two important centralities from NAPS network analysis: Betweenness and Closeness. Betweenness centrality measures the number of shortest paths passing through a node within a network. High betweenness nodes, also called bottlenecks, monitor the flow of information within a network. Whereas closeness centrality calculates the average distance of all the shortest paths between a node and every other node within a network. It measures which nodes are efficient in communications with the rest of the network. High closeness should indicate the proximity of a node to all other nodes. Nodes with high closeness are also initiators of effective allosteric communications.

**Figure 5:**
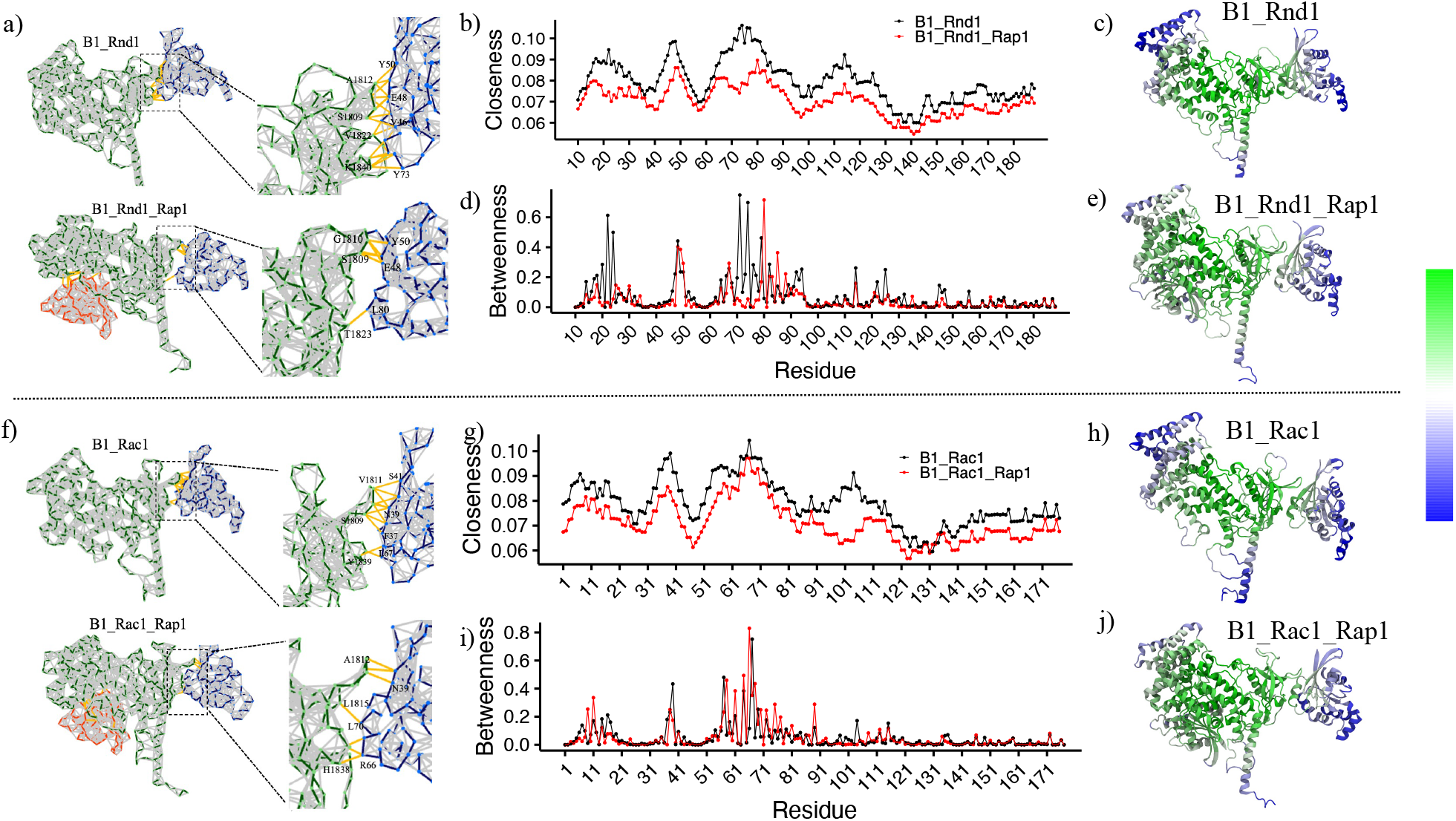
Network analysis of major cluster conformation. a) Rnd1 complexes, where zoom-in view shows few interacting residues b) Closeness centrality vs Residue for Rnd1 complexes, where B1_Rnd1 is colored in black and B1_Rnd1_Rap1 is colored in red. c) Closeness centrality values color-coded on the B1_Rnd1 complex. d) Betweenness centrality vs Residue for Rnd1 complexes, where B1_Rnd1 is colored in black, B1_Rnd1_Rap1 is colored in red. e) Closeness centrality values are coded on the B1_Rnd1_Rap1 complex. f) Network view of Rac1 complexes, where zoom-in view shows few interacting residues g) Closeness centrality vs Residue for Rac1 complexes, where B1_Rac1 is colored in black and B1_Rac1_Rap1 is colored in red. h) Closeness centrality values color coded on the B1_Rac1 complex i) Betweenness centrality for Rac1 complexes, where B1_Rac1 is colored in black, B1_Rac1_Rap1 is colored in red with the function of residues. j) Closeness centrality values color-coded on B1_Rac1_Rap1 complex.

**Figures 5b** and **5d** show closeness and betweenness values for all residues in Rnd1 with and without Rap1b. **Figures 5g** and **5i** show the same for the Rac1 systems. We observed that closeness values for B1_Rnd1 complex are significantly higher than B1_Rnd1_Rap1 complex, which suggests that in B1_Rnd1 complex, residues are more effectively communicating, suggesting a tighter coupling of interactions compared to B1_Rnd1_Rap1 complex. Interestingly, residues that interact with Plexin-B1 (residues 40-50, 70-80), had more closeness in the B1_Rnd1 system than the B1_Rnd1_Rap1 system, also indicated on the protein structure, color-coded with closeness values (**Figure 5c and 5e**). We also observe slightly more betweenness for residues spanning from 70-80, which indicates that region 70-80 in Rnd1 may act as a bottleneck for proper information flow in Plexin-B1 interaction. For B1_Rac1, the region of residues 30-40, which overlaps the nucleotide conformationally sensitive Switch I region of his GTPase and an interacting region for Plexin-B1, is shown to have higher values of closeness, but we did not observe many changes in betweenness values.

Overall, these results suggest that in the absence of Rap1b, the B1_Rnd1 protein complex is more stable with more connections. However, this effect was not observed for the Rac1 GTPase, indicating that Rap1b has a differential impact on the extent and strength of the interaction network between plexin and the GTPases.

### 2.4 Difference in Rap1b Dynamics in the presence of Rnd1 vs Rac1

Plexin has been identified as possessing an enzymatic function that activates Ras proteins, such as Rap1 and R-Ras, by hydrolyzing their GTP. This function classifies Plexin as a GTPase-Activating Protein (GAP), with a specific GAP domain dedicated to this catalytic activity. Therefore, understanding how interactions with Rap1b are influenced by interactions with various Rho-GTPases is crucial. Next, we examined how interactions between Rnd1 and Rac1 with the Plexin-B1 RBD affect the interactions between the Plexin-B1 GAP domain and Rap1b.

To investigate, we analyzed three simulations: B1_Rap1, B1_Rnd1_Rap1, and B1_Rac1_Rap1. We plotted root mean square fluctuations (RMSF) for Rap1b (residues 2–167) across these complexes (**Supplemental Figure 6**). No significant RMSF changes were observed, except for the B1_Rnd1_Rap1b system, which displayed slightly elevated fluctuations in specific regions. We then performed local frustration analysis using the Frustratometer R package, which enables us to identify how energy is distributed in protein structures. The highly frustrated regions often indicate biologically important regions, whereas minimally frustrated regions often constitute stable folding regions. Local frustration analysis revealed significant differences in Rap1b. **Figures 6b, 6d**, and **6f** illustrate local frustration in Rap1b residues within the B1_Rap1, B1_Rnd1_Rap1, and B1_Rac1_Rap1 systems, respectively. Among these systems, B1_Rac1_Rap1b exhibited minimal frustration (colored green), indicating limited large-scale conformational changes in Rap1b. Conversely, in the B1_Rnd1_Rap1 complex, Rap1b showed regions enriched with high-frustration interactions (colored red).

**Figure 6:**
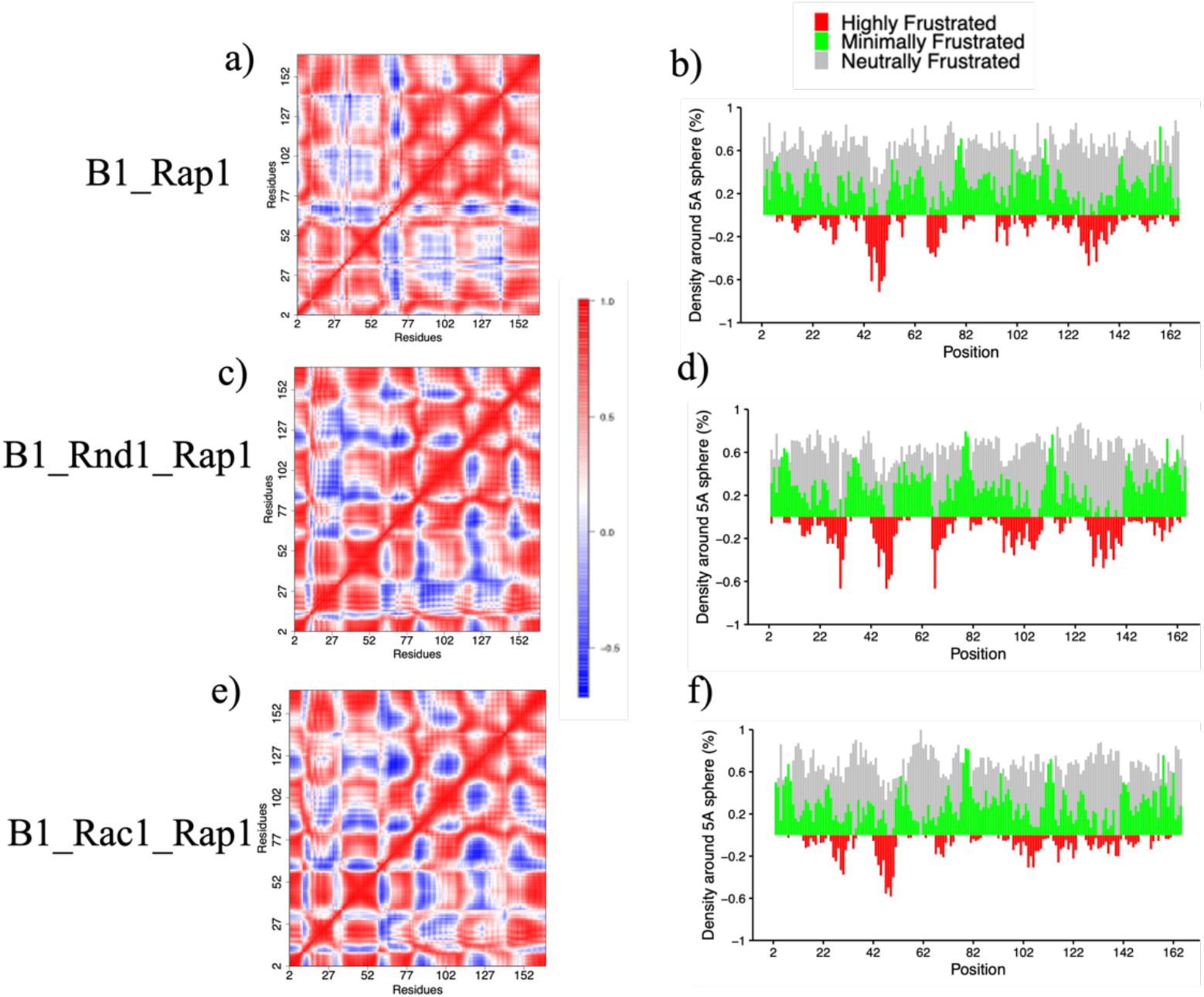
a), c), and e) Dynamical cross-correlation analysis for the Rap1b protein in all three complexes, where positive values indicate correlation and negative values indicate anti-correlation. b), d), and f) Local frustration on Rap1b protein structure for three systems.

We then assessed the extent to which these atomic fluctuations are correlated in these three systems by calculating the dynamical cross-correlation matrix for Rap1b using the Bio3d package (dccm function), as shown in **Figure 6**. The DCCM analysis identifies correlated motion between residues, also providing insights into protein flexibility and allosteric communication. We combine three replicas for each simulation using 50-200 ns simulation time for this calculation. We observe significant changes in Rap1b amino-acids motions when it is bound with Rnd1/Rac1 and when it is unbound. Our results indicate that introducing Rnd1 or Rac1 leads to changes in correlated atomic motions. Specifically, the B1_Rap1b system demonstrated more positively correlated motions, especially between residues 76-140. This difference was more significant in B1_Rnd1_Rap1 than in B1_Rac1_Rap1. Interestingly, this region (residues:76-140) was also enriched with high frustration interactions in the B1_Rnd1_Rap1 system. These differences suggest that the binding of Rho-GTPases to the RBD region may influence the interactions between Rap1b and Plexin-B1, specifically motions within Rap1b are altered depending on the type of Rho-GTPases (**Figure 6**). Our analysis also suggests that Rap1b itself is more stable when it is in complex with Rac1 (**Figure 6f**).

#### Network dynamics changes in Rap1b

In a similar manner to Rho-GTPases, we also performed network dynamics focusing on Rap1b interactions with Plexin-B1. Here, we plotted closeness (**Figure 7a**) and betweenness (**Figure 7b**) centralities for Rap1b residues for all three systems. The closeness values were significantly higher in B1_Rap1 compared with B1_Rnd1_Rap1/B1_Rac1_Rap1 which indicates that amino acids in the B1_Rap1 system have more proximity to other residues resulting in effective communication and connection. This is similar to our findings at the RBD-Rac1/Rnd1 end where the distant binding of Rap1b also reduced network closeness. We visualized the network connection between Rap1b and Plexin-B1 in **Figures 7c** and **7d** and observed that in B1_Rac1/Rnd1_Rap1(∼ 11-14 connections) there are a similar number of connections compared to B1_Rap1(∼14 connections), but with the different closeness among Rap1b residues. These findings suggest that Plexin-B1 complexed with Rnd1/Rac1 in the presence of Rap1b, alters the interaction and dynamics of the GAP domain and Rap1b itself. These trends are also apparent when the closeness and centrality are grouped for the different RBD and plexin-B1 complexes, examining their averages (**Supplementary Figure 7**).

**Figure 7:**
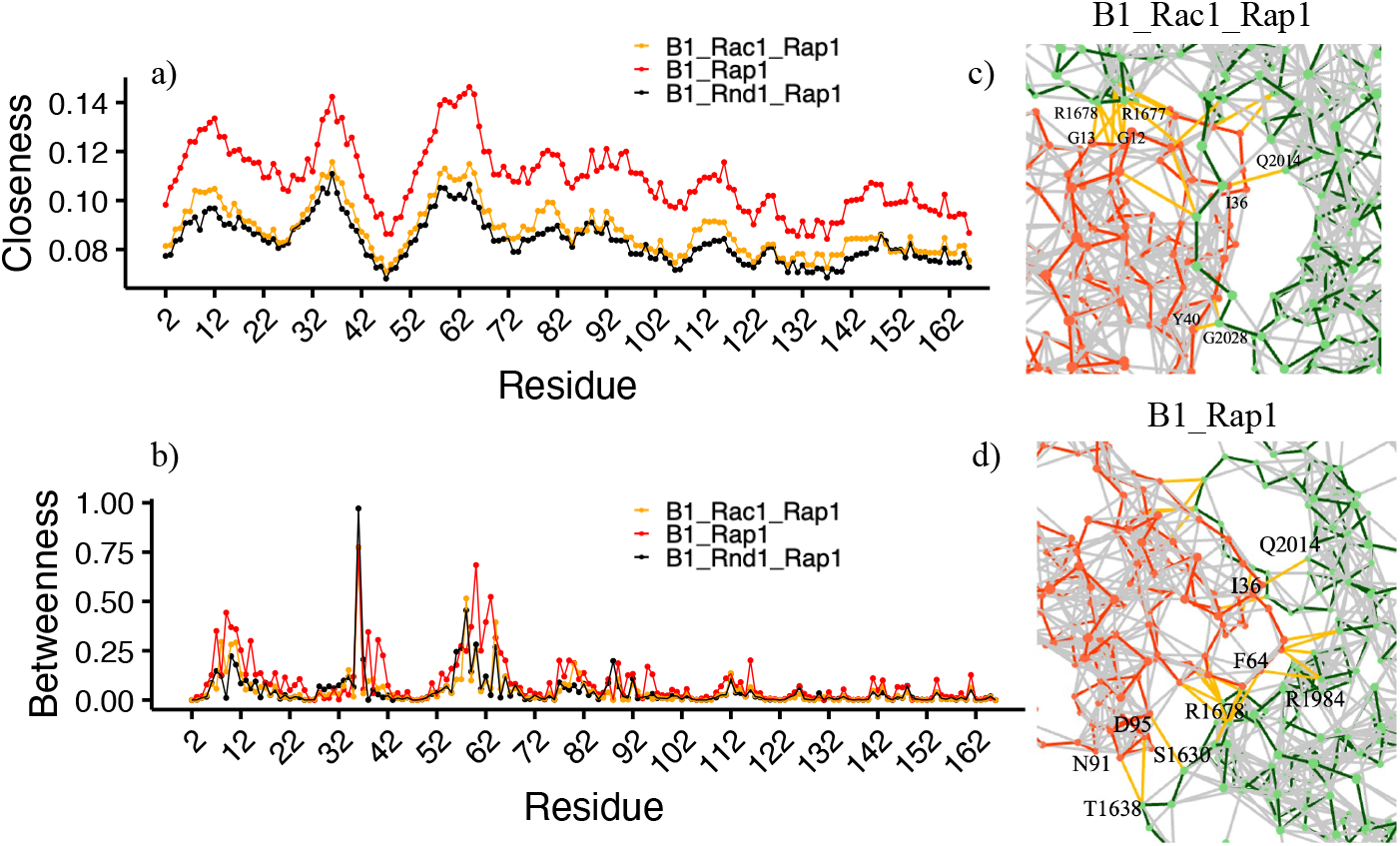
Network centralities vs Residue a) Closeness b) Betweenness where B1_Rac1_Rap1 is colored in orange, B1_Rnd1_Rap1 in black, and B1_Rap1 in red. c-d) Snapshot of network connections between Plexin-B1(green) and Rap1b (red).

### 2.5 Analysis of local protein dynamics/stability by hydrogen-deuterium exchange

We carried out two sets of HDX-MS experiments. First, plexin-B1 ICR with and without Rac1 and Rnd1 bound, as well as with Rap1b bound (without Rac1/Rnd1), and then, a few years later, second, of the trimeric plexin:Rac1/Rnd1:Rap1b complex (see Materials and Methods). The concentration of plexin was a limiting factor as there were two protein dilution steps before sample processing by quench and digestion into peptides in the instrument. Given measured Kd’s for the RBD: Rac1 and RBD: Rnd1 interaction of 2.5 ∼4 μM, we estimate that > 60% of plexin is in the bound state while undergoing amide hydrogen to deuterium exchange. The binding affinity of Rap1b is similar. However, when in the ternary mixture in D_2_O/H_2_O buffer, the concentration of plexin is 50% less than in the binary mixture but with GTPase concentrations are similar to the first set of experiments. This means that the expected amount of plexin bound is ∼ 40%. Although this fractional binding will likely reduce the magnitude of deuterium exchange, the off-rate is significant and there will be a difference in an exchange between bound and unbound plexin, as is evident from the data, because the bound plexin features only one population of hydrogen/deuterium occupancy, indicative of fast protein dynamics.

Data analysis and curation are described in Methods and in the Supplemental Methods, including **Supplemental Figures 8, 9, 11-13 and Supplemental Table 1**. Before data curation, the yield was 128 peptides at 91.2% coverage (3.6 redundancy) whereas the newer system processed 327 peptides at the same coverage (7.2 redundancy), a testament to the increased sensitivity and resolution. Because of the higher concentrations of the GTPases (10-20x excess), the chromatogram shows some overlap and after curation 185 peptides can be used (85.3% coverage, 3.9 redundancy). The results of deuterium fractional uptake are plotted as a function of the plexin ICR sequence in Fig. 8 for B1_Rac1 and B1_Rnd1 compared to plexin-B1 by itself.

**Figure 8.**
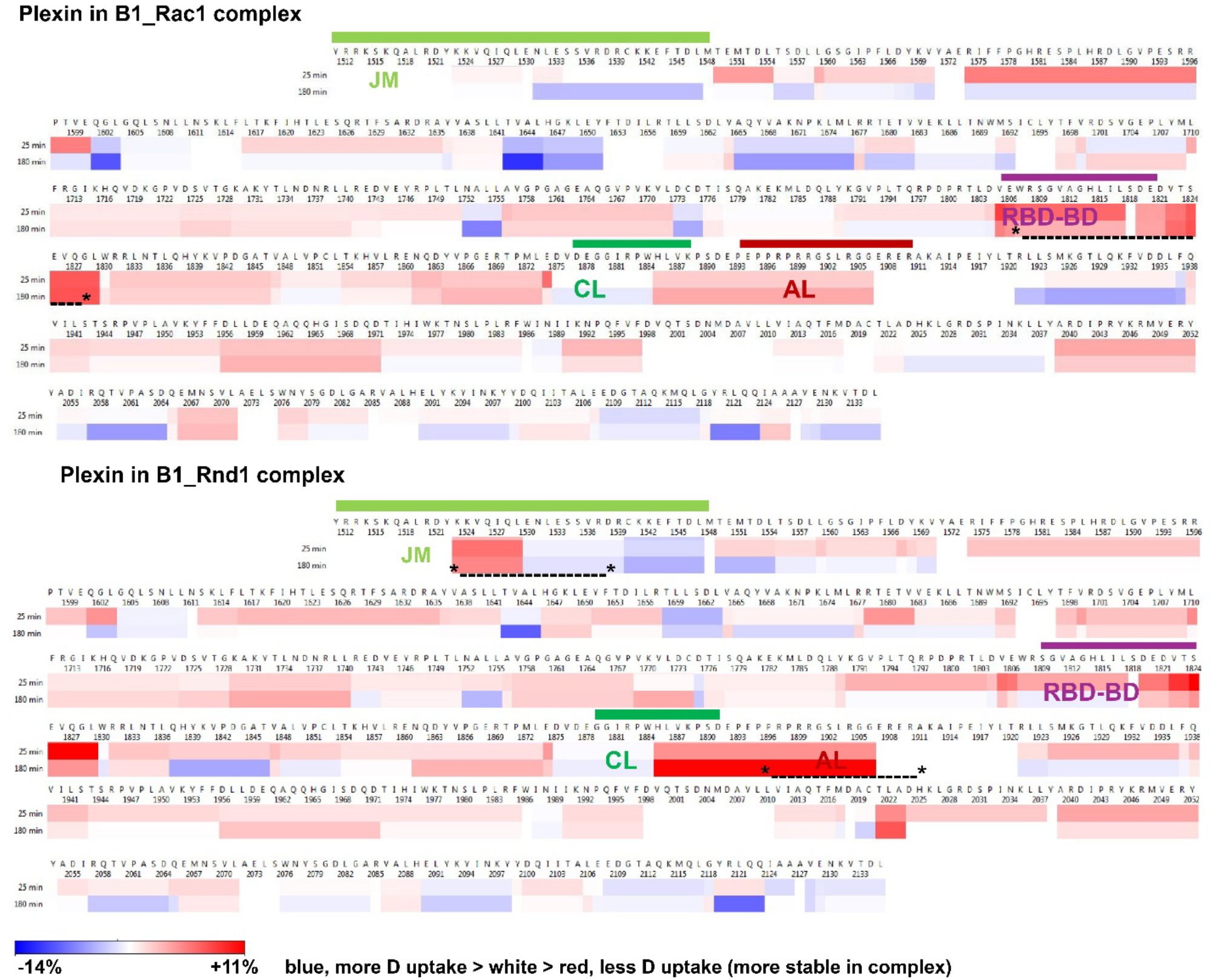
Protection of amides at binding interfaces: >110 protected peptides include most of the JM, RBD-binding and dimerization, coupling loops, and the RBD-GAP interface regions. We compared deuterium (D) uptake in plexin-B1 after 25 & 180 min, vs. its uptake in Rac1:plexin and Rnd1:plexin complexes, at pH 7.0, 25 oC. The difference data between bound-unbound is shown, demonstrating that the RBD-binding domain, but also the JM and activation switch loop, are more stable (incorporate less D, red; esp. in the Rnd1 complex. (indicated by * *, respectively). The coupling and Activation loop, as well as the RBD position, are indicated above the sequence by green, red, and violet, respectively.

The HDX data in **Figure 8**, show that in the case of Rnd1 binding the JM region, time points 25 and 180 mins (other data not shown) have greater protection than with Rac1 bound. This is also true for peptides that partially overlap the coupling and activation loop as well as a peptide res. 2024-2037, which precedes the region in the GAP domain known to bind substrate Ras GTPases, suggesting that this region is more persistently structured with Rnd1 bound to the RBD, even though it is spatially distant and is not bound. A second region in the GAP domain (res. 1685-1675), which just precedes the segment that contains the two catalytically active Arg (R1677 and 1678) to help Rap1b with GTP hydrolysis is more flexible with Rac1 bound to the RBD than in unbound plexin, whereas amides are more protected when Rnd1 is bound to the RBD. By contrast, the RBD is slightly more protected when Rac1 is bound. Thus, there is a wide range of differences in HDX behavior in different regions, but notably in regions, with the exception of the JM, known to bind either Rho or Ras GTPases.

In examining the differences between HDX in plexin:Rac1:Rap1b: and plexin:Rnd1:Rap1b complexes (**Supplemental Figure 10**) it is striking that relative to plexin ICR on its own, differences are mostly in the direction of greater protection, but less strong than seen in the plexin:Rac1/Rnd1 complexes above. Overall, the differences seen in the plexin:Rac1/Rnd1 complexes are greatly diminished when Rap1b is bound. Two regions that stand out as having slightly greater protection in the complex with Rap1b, than in the unbound plexin ICR are 1848-1875 and 1958-1986, but these are not much different between Rac1/Rnd1 complexes, which was already indicated in the absence of Rap1b.

## 3. DISCUSSION

The interactions of Plexin-B1 with Rho-GTPases (Rnd1 and Rac1) and Ras-GTPase (Rap1b) play a crucial role in regulating plexin signaling pathways in cells. A few X-ray crystal studies have suggested a role of Plexin-GTPase interactions, through structural changes at the Rho-GTPase Binding Domain (RBD)^19,28^ and its dimerization loop^11,19,^ and while other studies focused on charges in the juxtamembrane region, which can attached to the GTPase Activating Protein (GAP) domain^22,36^. In this study, we explored how bound Rnd1 and Rac1 influence Rap1b dynamics and, conversely, how bound Rap1b influences Rho-GTPase dynamics (Rnd1/Rac1). The molecular dynamics (MD) simulations and hydrogen-deuterium exchange mass spectrometry (HDX-MS) also report on the overall stability and structural behavior of the Plexin-B1 GTPase complexes. For MD simulations, we ran a total of six systems, each with three replicas of 200 ns in the solution: Plexin-B1 (B1_only), Plexin-B1 with Rnd1 (B1_Rnd1), Plexin-B1 with Rac1 (B1_Rac1), Plexin-B1 with Rap1 (B1_Rap1), Plexin-B1 with Rnd1 and Rap1 (B1_Rnd1_Rap1), and Plexin-B1 with Rac1 and Rap1 (B1_Rac1_Rap1).

We analyzed four systems with and without Rnd1 and Rac1 bound to plexin-B1 to assess how Rnd1 and Rac1 dynamics with plexin-B1 differ in the presence and absence of Rap1b. Examining the average root mean square fluctuations (RMSF) of each Rho-GTPase, we found that Rnd1 showed slightly higher fluctuations, particularly in residues 90–100, which are near the RBD region. Additionally, interaction energy analysis indicated that the B1_Rnd1 system was slightly more stable than the B1_Rnd1_Rap1 system. However, this stability trend did not hold for Rac1; the B1_Rac1 system was less stable than B1_Rac1_Rap1, suggesting that Rap1b affects Rnd1 and Rac1 differently. Network analysis showed that in the absence of Rap1b, the B1_Rnd1 complex was more stable, displaying more connections and greater closeness among residues. This effect was also observed for Rac1 but is not significant, indicating that Rap1b influences the interaction network between plexin-B1 and the GTPases in distinct ways.

Our findings suggest that plexin-B1-RBD interactions with Rnd1 are more favorable in the absence of Rap1b. In contrast, although the B1_Rac1 system displayed slightly more connections than the B1_Rac1_Rap1 complex, these connections were less stable, suggesting that the effect of Rap1b binding is more pronounced for Rac1. These findings corroborate earlier observations, detecting an isoform-specific dynamics of Rho-GTPases interacting with Plexin-B1.^28^ As shown in our MD simulations in this report as well as previously^26^, Rnd1, and Rac1 exhibit distinct structural dynamics when complexed with Plexin-B1, which –as shown here- are further modulated by Rap1b binding. Similar to the findings of Zhang and Buck (2017)^26^, Rnd1 demonstrates a larger number of dynamic network connections with Plexin-B1 in the absence of Rap1b, indicating a stronger and more stable interaction. Rac1’s interactions with Plexin-B1, while still stable, appear less robust, with the network analysis revealing fewer and less stable connections when compared to Rnd1.

Upon examining how Rnd1 and Rac1 binding influences Rap1b dynamics, our local frustration analysis of Rap1b revealed that the B1_Rac1_Rap1b complex showed minimal frustration, indicating limited large-scale conformational changes in Rap1b. In contrast, the B1_Rnd1_Rap1 complex displayed regions in Rap1b with high frustration, which were also positively correlated. These findings suggest that Rho-GTPase binding to the RBD region may affect interactions between Rap1b and Plexin-B1, specifically impacting Rap1b motions. Consequently, Rap1b dynamics vary depending on the specific Rho-GTPase involved. However, in terms of network connections between Rap1b and Plexin-B1, there were very few differences between Rnd1 and Rac1, with slightly more closeness with Rac1 bound Plexin-B1. These results suggest that Plexin-B1 bound with Rac1 is more favorable for Rap1b rather than when Rnd1 is bound. These results also correlate well with our previous observation in this study that Rap1b bound with Plexin-B1 is more stable with Rac1 . Consistent with previous studies of Ras-GTPase interactions with Plexins – the role of an activation switch loop, which is part of the relatively flexible segment linking RBD to the C-terminal GAP region (Li et al., 2021)^27^, our results also show that Rap1b binding alters the interaction network of Plexin-B1 and its associated Rho-GTPases. Specifically, the binding of Rap1b to Plexin-B1 reduces the stability of the Rnd1 complex but enhances the stability of the Rac1 complex. This suggests that Rap1b preferentially stabilizes Plexin-B1 interactions with Rac1 over Rnd1.

In Plexin-B1, we observed notable changes primarily in the RBD (Rho-GTPase binding domain) and the RBD-GAP connecting loop regions. Across the six systems, the RBD in the B1_Rac1_only system aligned with our observation that the B1_Rac1 system was less stable in the absence of Rap1b. Stabilization was also seen in regions 2013–2043, where the Rap1b binding region was more stable in the B1_Rac1_Rap1 system and remained unchanged for Rnd1 systems. Besides the RBD, the B1_Rac1 system also showed higher flexibility in the RBD-GAP connecting loop. When comparing principal components based on the positions of alpha carbon atoms, we observed that the B1_Rac1_only system transitioned between conformations more frequently than the other systems. This instability might reflect a greater requirement for cooperative stabilization when Rac1 is bound to Plexin-B1. Further inspection indicated that this increased flexibility may stem from the flexible nature of the plexin RBD position in the complex and of the RBD-GAP connector regions.

From HDX-MS studies, we found that the effects of Rho-GTPase binding on the regional flexibility of the complex were attenuated when Rap1b was bound to Rho-GTPases Rnd1/Rac1. This suggests that Rap1b-induced conformational changes in Plexin-B1 influence fluctuations and accessibility of the RBD, potentially modulating downstream signaling pathways^37^. This observation is consistent with our network analysis, where we noted more connections between plexin and GTPases in the absence of Rap1b, particularly for Rnd1 and to a lesser extent for Rac1. HDX-MS data also indicate that Rac1 and Rnd1 have different effects on plexin activation, though the molecular mechanism is just beginning to come into focus. Specifically, we are not able to pin-point from the present simulations which modulated dynamics in the presence of the Rnd1/Rac1 GTPases constitutes the critical difference that may lead to different functional behavior. Thus, to understand how these structural and network changes in plexin-GTPase interactions impact the overall function of the intracellular region of plexin-B1,further experimental studies are required. The simulation suggested networked interactions can now be explored by experimental site-directed mutagenesis. However, the incorporation of additional components such as the membrane and the dimeric structure of plexin is likely to be critical and we have planned this for our future research.

In summary, in this study, we used molecular dynamics simulations and HDX-MS to explore inter- and subdomain-level structural dynamics of Rac1 and Rnd1 in the presence and absence of Rap1b. Our findings revealed that Rho-GTPases Rnd1 and Rac1 exhibited differential effects on Plexin-B1 in the presence and absence of Rap1b. The interaction of Ras and Rho-GTPases with Plexin-B1 not only influences Rho-GTPase dynamics but also impacts the dynamics of both Rap1b and Plexin-B1.

## 4. MATERIALS AND METHODS

The initial structure of the Plexin-B1 intracellular region (ICR: 1513-2135) was taken from the Protein Data Bank (PDB ID: 3SU8)^12^, and missing residues were modeled using Modeler^38^. Following the Plexin-B1 structure, we began modeling the GTPases onto it. Structural superimposition was conducted for three GTPases: Rnd1, Rap1b, and Rac1, utilizing deposited structures of Plexin bound to GTPases in the Protein Data Bank (PDB ID: 2rex 3su8, and 4m8n)^12,39,40^. We prepared a total of six systems in solution: Plexin-B1 (B1_only), Plexin-B1 with Rnd1 (B1_Rnd1), Plexin-B1 with Rac1 (B1_Rac1), Plexin-B1 with Rap1 (B1_Rap1), Plexin-B1 with Rnd1 and Rap1 (B1_Rnd1_Rap1), and Plexin-B1 with Rac1 and Rap1 (B1_Rac1_Rap1). Systems were prepared using the *Charmm-gui* webserver: Solution builder plugin^41^. All the systems were solvated with TIP3 water molecules in a cubic box with 150 mM NaCl salt concentration, totaling ∼ 332,000 atoms. In addition, a GTP and an MG ion were included in each of the GTPases at the appropriate crystallographic binding site.

All-atom molecular dynamics simulations were conducted using *Gromacs 2023*^42^ and the CHARMM36m forcefield^43^. We followed the standard simulation protocol, where we initially performed energy minimization and equilibration steps^31,44,45^. After equilibration, production runs were carried out at constant pressure and temperature (NPT) with a 2 fs time step. Semi-isotropic Parrinello-Rahman pressure coupling was employed to maintain a pressure of 1 bar, while the LINCS algorithm^46^ was used to constrain bonds involving hydrogen atoms. Temperature was regulated using Nose-Hoover coupling^47^. Van der Waals and electrostatic interactions were truncated at 12 Å, and long-range electrostatics were handled using the Particle Mesh Ewald (PME)^48^ method with periodic boundary conditions. A representative initial configuration of the system is shown in **Figure 1**. Each simulation was run for 200 ns, and to ensure reproducibility and reliability, two additional 200 ns replicas were performed, totaling 600 ns for each system.

### Analysis of MD trajectories

Most analyses were performed using embedded modules in the Gromacs simulation package 2023. We analyzed our simulated trajectories for all replicas using *gmx_rms, gmx_rmsf, gmx_anaeig*, and *gmx_covar* tools of the Gromacs 2023, to extract root mean square deviation, root mean square fluctuations, principal component (PC) analysis, and covariance matrix, respectively. For PC analysis, we used the *gmx_covar* module which performs calculation and diagonalization of the covariance matrix and outputs the corresponding eigenvectors and eigenvalues based upon positional fluctuations of C_*α*_ atoms as described as *C*_*ij*_ = ⟨ (*x*_*i*_ − ⟨ *x*_*i*_⟩) (*x*_*ij*_ − ⟨ *x*_*j*_⟩)⟩ where x_i_/x_j_ is the coordinate of i/j_th_ atom and < > represents the ensemble average. The dynamic cross-correlation analysis for all the systems was computed using the *dccm* function of Bio3D^36^. This function calculates the covariance matrix based on mutual information between all Cα atoms in the interface structures. For this calculation, we combined each system’s last 50-200 ns of the replica trajectories. All the graphs were plotted using R (version 4.2.0). Visualizations were done using VMD (version 1.9.4) ^49^ and Pymol software.

### Network Analysis and Frustration Analysis

We conducted network analysis for all systems using the NAPS (Network Analysis of Protein Structures) webserver^34^ which allows interactive network visualization for protein-protein complexes and includes features for several network analysis centralities. The detailed protocol of the NAPS application is described by *Chakraborty et. al* (2016)^34^. It uses C*α* atoms as a network node and calculates several global properties. To perform network analysis, we first perform cluster analysis using *gmx_cluster* by combining trajectories from all replicas for each system (totaling 600 ns for each system). Subsequently, we utilized the protein conformation from the major cluster (representing ∼60-70% of the conformations) as input for network analysis. We then extracted the data files for network centralities (Closeness and Betweenness) and later graphed them. The two key centralities we focused on were closeness and betweenness centralities. Closeness centrality (CC) for a residue i (ranging from 0 to 1) measures how close that residue is to all other residues in the network. It highlights residues that are highly efficient in transmitting information throughout the network. In contrast, betweenness centrality (BC) reflects the importance of a residue in controlling the flow of communication. Residues with higher BC values lie on critical communication paths and may serve as strategic targets in drug discovery. BC evaluates the shortest paths between residues j and k that pass through residue iii, assigning a value to iii. CC, however, calculates the distance from residue i to all other residues, assigning a value to residue i ^50^.

We conducted frustration analysis on simulated protein structures across all complexes using the Frustratometer-R-package^51^, an algorithm that is inspired by energy landscape theory. This analysis provides valuable insights for identifying and quantifying regions of local frustration within proteins. Regions with significant conformational changes often show clusters of highly frustrated interactions, helping pinpoint areas critical to protein function^44^. To explore the free energy landscape of plexin-GTPases, we performed cluster analysis and selected the major conformation for each system. We then use this major conformation structure as an input for frustration analysis.

### HDX-MS

GTPases and Plexin-B1 ICR were expressed in and purified from *E. Coli* as described in a previous paper^27,52^. In the case of Rac1 constitutively active mutant Q61L was used and in the case of Rap1b, the protein was loaded with GMPPNP to slow GTP hydrolysis. GTP hydrolysis in Rnd1 is already very slow. The stock solutions were at 110 μM plexin-B1 and 120 μM Rap1b, 320 or 530 μM Rnd1, and 180 or 440 μM Rac1.Q61L. All proteins were in buffer E (see below) flash frozen and stored at −80 °C until use. Buffers used were as follows: Equilibration buffer (E) 20 mM Tris.HCl pH 7.5, 150 mM NaCl, 5% glycerol, 2 mM TCEP, 0.005% Tween-20, 4 mM MgCl_2_ in H_2_O. Labeling buffer (L) 20 mM Tris.HCl pH 7.1 (pD 7.5), 150 mM NaCl, 5% glycerol, 2 mM TCEP, 0.005% Tween-20, 4 mM MgCl_2_ in D_2_O. Quench buffer (Q) 100 mM Tris.HCl pH 2.4, 2.0 M Gdn.HCl, 0.5 M TCEP. Plexin was initially diluted to 30 μM in buffer E, then either it was further diluted to 7.5 μM in E buffer (plexin alone sample) or mixed 1:1 with Rac1 or Rnd1 (10 μL each) and then combined with 20 μL Rap1b (final concentration 7.5 μM) arriving at final mixture of 1:1:2 Plexin/ Rac1 or Rnd1 / Rap1b. In the case that no Rap1b is added the concentrations are 2-fold higher. The concentrations of proteins during labeling were estimated for plexin: Rac1:Rap1b as molar excess of Rac1 and Rap1b as ∼ 1: 14.5: 8.3 and for plexin: Rnd1:Rap1b as molar excess of Rnd1 and Rap1b as ∼ 1: 21.4: 8.3. The complexes were incubated at room temperature for 1 min before labeling. Conditions were reference (E) 10 sec, 25 min, and 180 min. For the experiments without Rap1b, also a sample as taken after 2.5 min of exchange. Quenching was done by addition of 60 μL of buffer Q, waiting 3 mins at 0°C. The total dilution factor of the plexin was 30-fold giving with a 100 μL loop a protein (plexin) load of 25 pmol. This was injected into an Enzymate BEH pepsin column followed by a Vanguard BEH C18 column and in-line transfer into the Waters Inc. Acquity UPLC M-class instrument with the HDX manager system. For the experiments without Rap1b, which occurred several years prior, but with the same protocol an older system, Synapt G2-Si system was used. Additional technical information on the method and analysis is available in the supplemental materials, including a figure showing the quality of representative data, a butterfly plot showing deuterium uptake, and a table of the Plexin:Rap1b data (**Supplemental Figures 8, 9, and Table S1**).

## 5. SUPPLEMENTAL MATERIALS

Additional molecular simulation data analyses including root mean square deviation, frustration of protein complexes, Interaction energy, HDX-LC-MS full protocol

## ACKNOWLEDGMENTS

We thank Gabrielys Encarnacion Lopez, Pravesh Shrestha, Maria Iannucci, and Jeannine Müller-Greven for help with protein expression and purification. NIH grants R01AG089561 and previously R01EY029169 support M.B. and the group. N.B. is funded by a postdoctoral fellowship NCI T32 (T32CA094186). We want to thank CWRU HPC for providing computing resources.

## Notes

### Competing Interest Statement

The authors have declared no competing interest.

### Summary of Updates

Section 2.5 has been updated. Some other sections have been arranged. A few supplemental figures added

