## Supplemental Information for "Computation model predicts Rho GTPase function with the Plexin Transmembrane receptor GAP activity on Rap1b via dynamic allosteric changes"

**Supplemental Figure 1:** Average root mean square deviation as a function of simulation time for each domain Plexin-B1, Rac1, Rnd1, and Rap1, within complexes, except for B1_only.

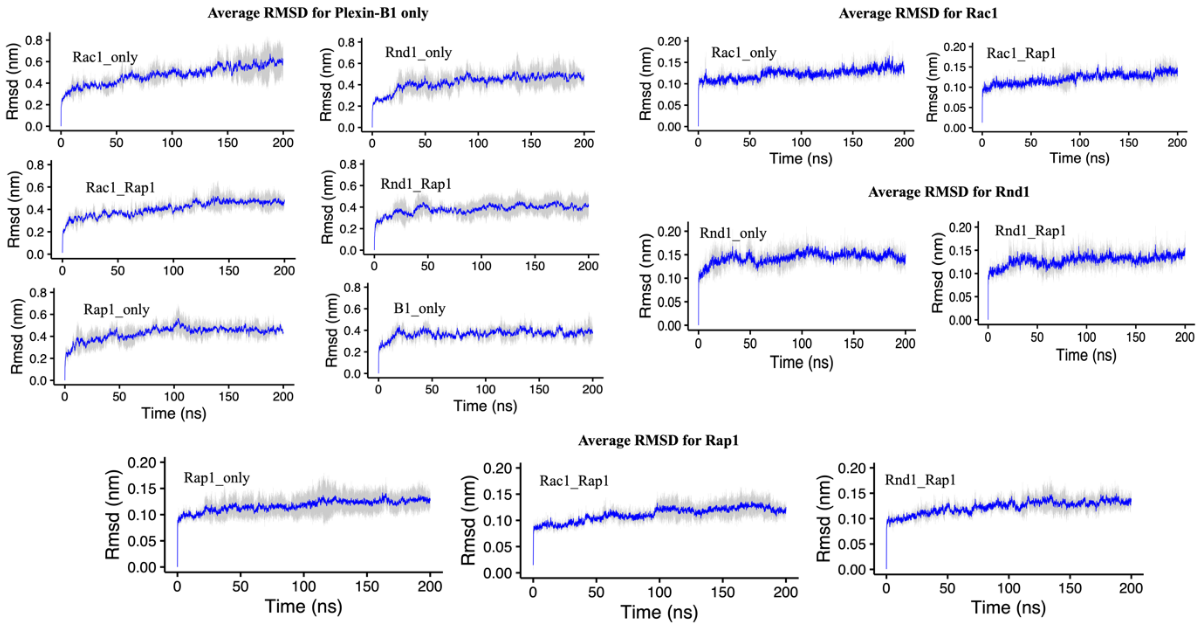


**Supplemental Figure 2:** Local frustration plotted for three regions in Plexin-B1: RBD region (residues: 1746-1852), one of the Rap1b binding regions (residues: 2013-2043), and RBD-GAP connecting loop (residues:1853-1913) in Plexin-B1 for all six systems, where highly frustrated regions are colored red, neutral in gray and minimally frustrated in green.

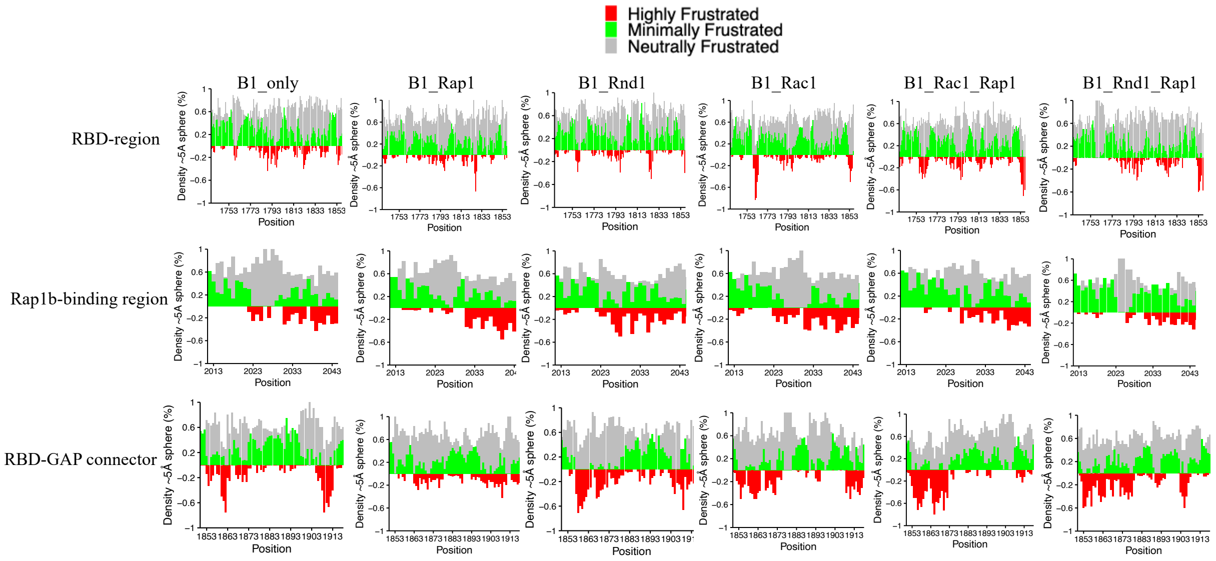

**Supplemental Figure 3:** Average Interaction energy from all replicas between Plexin-B1 and GTPases. The title of the plot shows the type of energy being calculated between respective protein structures.

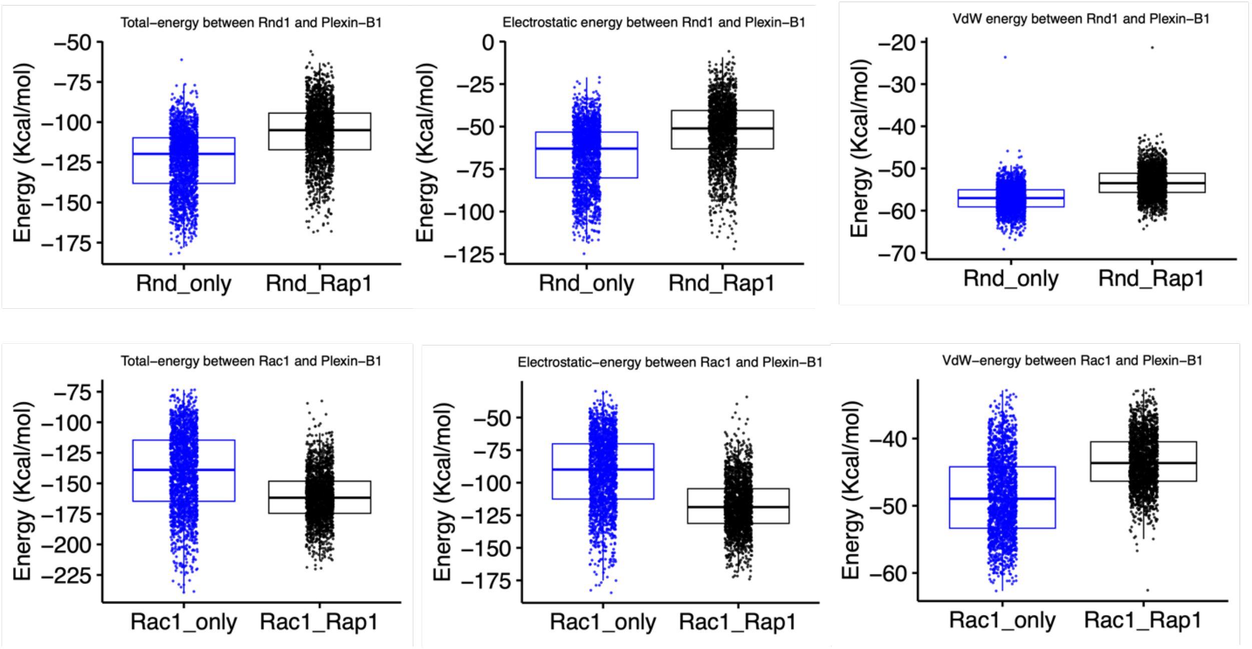


**Supplemental Figure 4:** Contact map from major conformation between Plexins and GTPases.

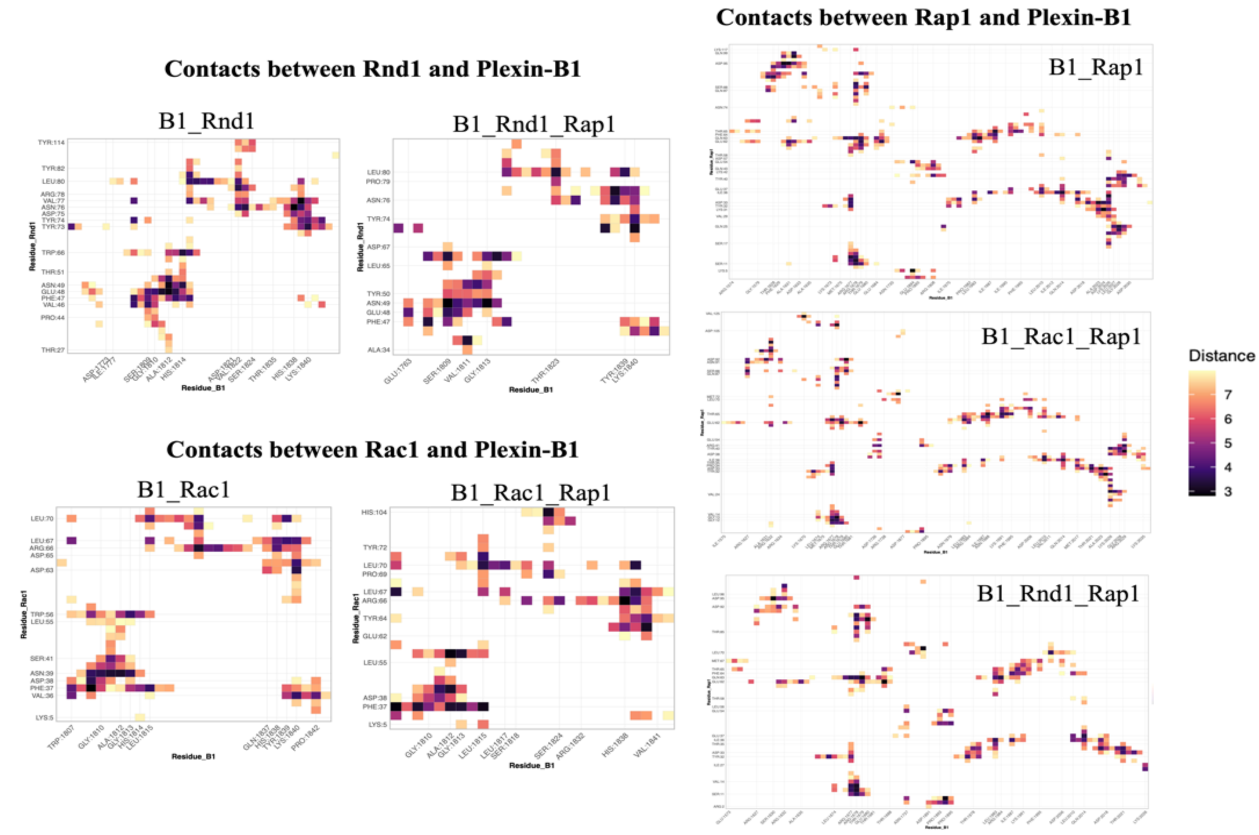

**Supplemental Figure 5:** Dynamical network analysis generated using NAMD/VMD software for four complexes, combining trajectories from all replicas.

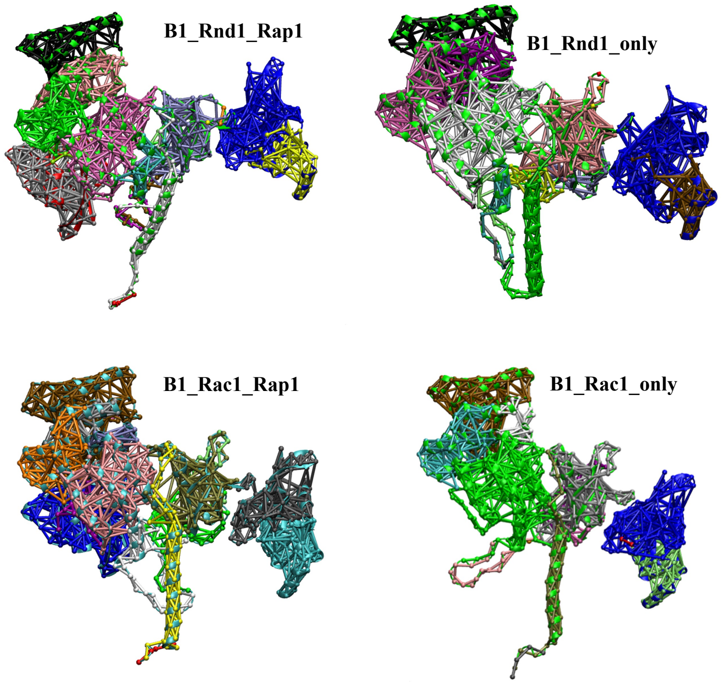


**Supplemental Figure 6:** Root mean square fluctuation vs Residue in Rap1b, where B1_Rac1_Rap1 is colored in orange, B1_Rnd1_Rap1 in black, and B1_Rap1 in red.

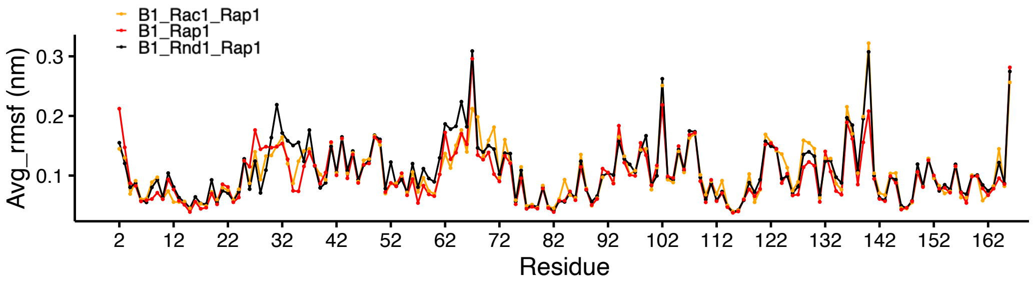

**Supplemental Figure 7:** Closeness Centrality vs Residue for Plexin-B1 and RBD only generated from network analysis

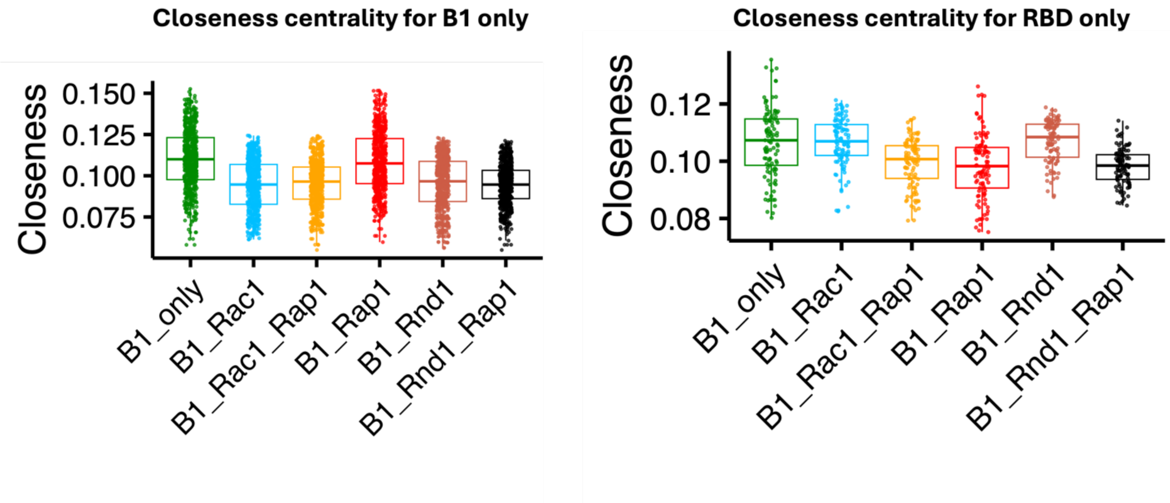


### Supplemental Methods: Full HDX LC-MS Protocol

A Waters ACQUITY M-Class system with HDX configured for online digestion (consisting of µBSM, ASM, and HDX Manager modules), coupled to a Waters SELECT Series Cyclic IMS mass spectrometer, was used for all HDX experiments, except for experiments without Rap1b, for which the same inlet was used but with a Waters Synapt G2-Si mass spectrometer. All injections were performed manually using a 250 uL gastight syringe (Hamilton). All HDX conditions were acquired in triplicates.

Samples (25 pmol plexin) were loaded onto an Enzymate BEH Pepsin Column (2.1 x 30 mm, 5 µm) kept at 15°C using water with 0.05% formic acid at a flow rate of 100-200 µL/min (see Gradient tables below) and online digested for 3 min, with resulting peptides being trapped and desalted on a VanGuard BEH C18 trap column (2.1 x 50 mm, 1.7 µm) kept at 0.2°C. After trapping/desalting, peptides were eluted from the trap column to an analytical ACQUITY UPLC BEH C18 column (1.0 x 100 mm, 1.7 µm) also kept at 0.2°C using a 12-min analytical gradient of water and acetonitrile, both with 0.1% formic acid, at 40 µL/min (see Gradient tables below) and directed to the mass spectrometer for detection, for a total of 15 min per injection.

**Analytical 12-min Gradient**

| Time (min) | Flow Rate (µL/min) | %A | %B |
|---|---|---|---|
| Initial | 40.0 | 95.0 | 5.0 |
| 7.0 | 40.0 | 65.0 | 35.0 |
| 7.5 | 40.0 | 15.0 | 85.0 |
| 9.0 | 40.0 | 15.0 | 85.0 |
| 9.1 | 40.0 | 95.0 | 5.0 |
| 12.0 | 40.0 | 95.0 | 5.0 |

**Trapping 3-min**

| Time (min) | ASM A1 Flow Rate (µL/min) |
|---|---|
| Initial | 100.0 |
| 1.0 | 200.0 |
| 2.2 | 200.0 |
| 2.5 | 100.0 |
| 3.0 | 100.0 |

Both Cyclic IMS and Synapt G2-Si instruments were operated in mobility data-independent acquisition (DIA) mode (HDMS$^{E}$) for data collection. Relevant tuning parameters for both are listed below. Parameters not mentioned were left at their default setting.

### Synapt G2-Si Parameters

| Tune Page | |
|---|---|
| Capillary Voltage | 2.9 kV |
| Cone Voltage | 25 V |
| Source Offset | 25 V |
| Source Temperature | 80ºC |
| Cone Gas | 0 L/h |
| Desolvation Temp. | 175ºC |
| Desolvation Gas | 400 L/h |
| Quad Profile | 400/500/600 (25%) |

| MS Method | |
|---|---|
| Acquisition Mode | HDMS$^{E}$ |
| Polarity | Positive (ESI+) |
| Analyser Mode | V |
| Data Format | Continuum |
| Scan Time | 0.4 s |
| Mass Range | *m/z* 50-2000 |
| HE CE Ramp | 20-40 V |
| Lockmass | GFP *m/z* 785.8426 |

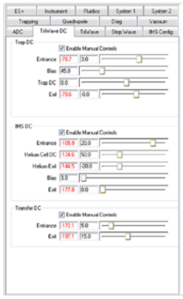

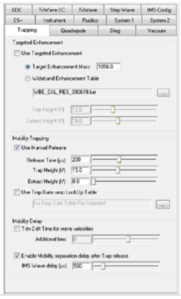

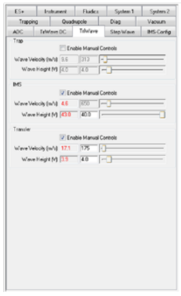

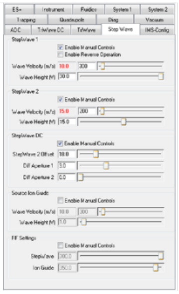

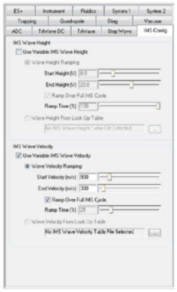

### SELECT Series Cyclic IMS Parameters

| Tune Page | |
|---|---|
| Capillary Voltage | 2.5 kV |
| Cone Voltage | 30 V |
| Source Offset | 10 V |
| Source Temperature | 100ºC |
| Cone Gas | 0 L/h |
| Desolvation Temp. | 450ºC |
| Desolvation Gas | 600 L/h |
| Nebuliser | 6 bar |
| Quad Profile | 400/500/600 (25%) |

| MS Method | |
|---|---|
| Acquisition Mode | HDMS$^{E}$ |
| Polarity | Positive (ESI+) |
| Analyser Mode | V |
| Data Format | Continuum |
| Scan Time | 0.4 s |
| Mass Range | *m/z* 50-2000 |
| HE CE Ramp | 20-50 V |
| Lockmass | GFP *m/z* 785.8426 |

| Cyclic Sequence Settings | |
|---|---|
| ADC Delay | Manual (Calculate) |
| Sideways Wave Velocity | 375 m/s |
| Static Wave Height | 23 V |
| Pusher per Bin | 2 |
| Passes | 1 (2 ms Separate) |

After data collection, reference RAW files were initially searched in ProteinLynx Global Server (PLGS) v.3.0.3 to generate peptide mapping references. A sequence databank was created using the clipped sequence of human Plexin-B1 from residues 1511-2135. Processing parameters were created with LE 250, HE 50 and GFP as doubly charged Lockmass (785.8426). For Workflow Parameters, the Plexin-B1 databank was selected with the following settings: Min. Fragment Ions Matched Per Peptide 3, Min. Fragment Ions Matched per Protein 7, Min. Peptides Matched per Protein 1, and Primary Digest Reagent set as Non-specific (for Pepsin).

Triplicates of ion accounting (IA) reference files were then loaded into DynamX v.3.0.0 and further filtered using parameters described earlier[1], see picture. All reference and labelling data were then processed in DynamX using an LE threshold of 250 to obtain deuterium uptake measurements and graphs, and data for all peptides was manually curated for consistency.

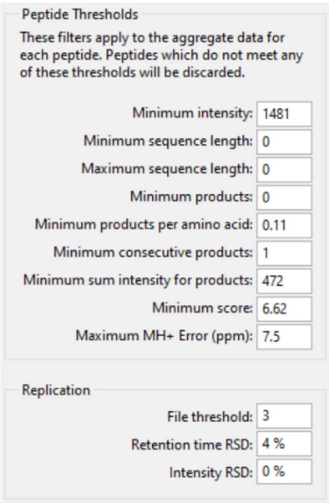


**Supplemental Figure 8. Figure Showing Data Quality**

HDX LC-MS data quality examples collected on the Cyclic IMS instrument for the ternary complex experiments. **A)** Stacked triplicate injection chromatograms of Plexin-Rnd1-Rap1b references (no labelling) at 25 pmol loads showing great reproducibility. **B)** Example spectra at 6.36 min for the same sample, demonstrating the complexity observed with typical HDX data. The insert shows a typical isotope profile measured with ~76,000 full-width at half maximum (FWHM) mass resolution. **C)** Spectral cleanup using the ion mobility separation performed on-the-fly during data processing with the DynamX software for peptide PCLTKHVLRENQ (1851-1862). The spectrum on the left shows the raw data and the one on the right shows the post-mobility cleanup result, greatly facilitating annotation and measurement of the deuterium uptake.

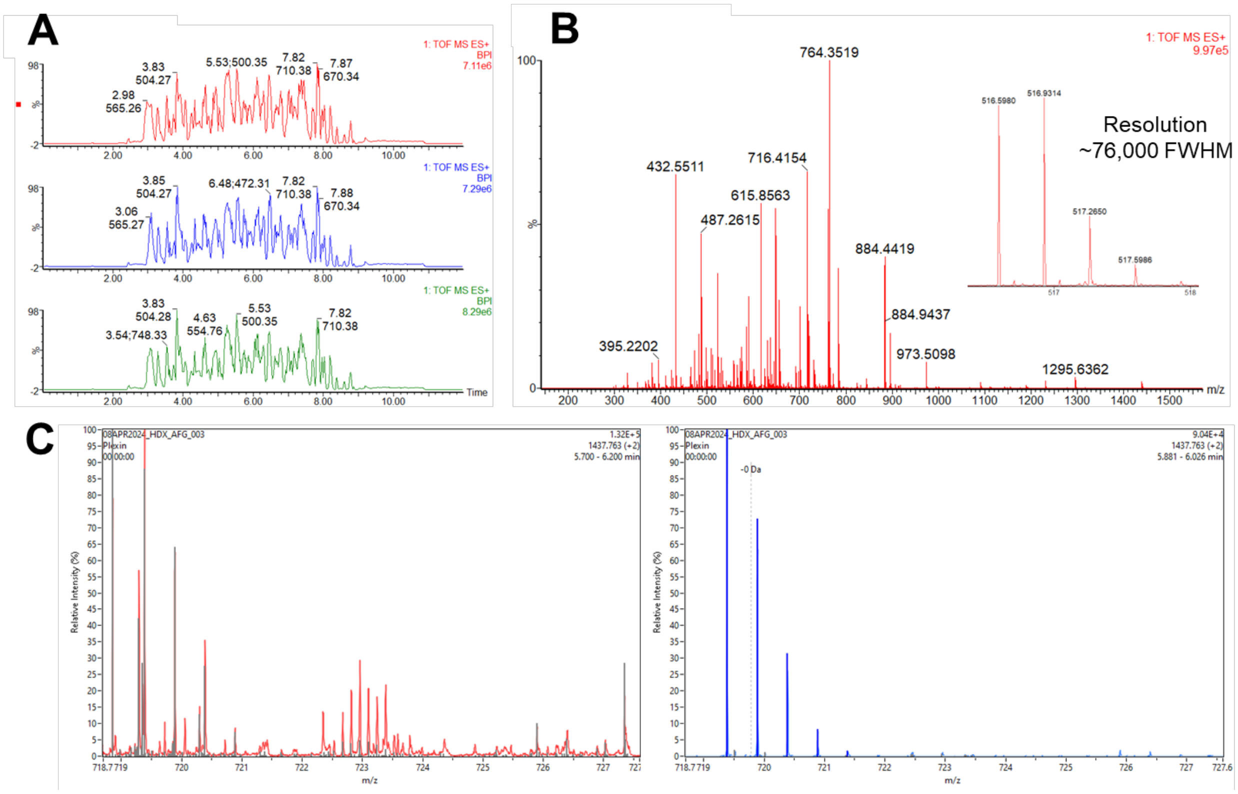

Norms for conducting and analyzing HDX-MS experiments have just recently been established[2,3] and were not well followed back in 2018 when the first dataset was acquired. These are demanding standards which were not available to us, given the limited access to the HDX-MS instrument at Waters, Inc., and are only slowly being adopted in the field. Presently, a difference in deuterium uptake beyond 0.2 to 0.7 Da is considered as significant (e.g.[4]). Here this is the case of the GTPase binding region in the RBD, and section preceding the Coupling loop (Fig. S9). A table of the Plexin:Rap1b data indicates standard deviations of degenerate peptides within the 2023 experiments, as less than 0.2 Da  (Table S1).

**Supplemental Figure 9.** Difference in deuterium update comparing Plexin with Plexin:Rac1 and Plexin:Rnd1 complexes, a) and b, respectively, plotted as a function of plexin-B1 intracellular region sequence. The difference colored lines indicate the timepoints

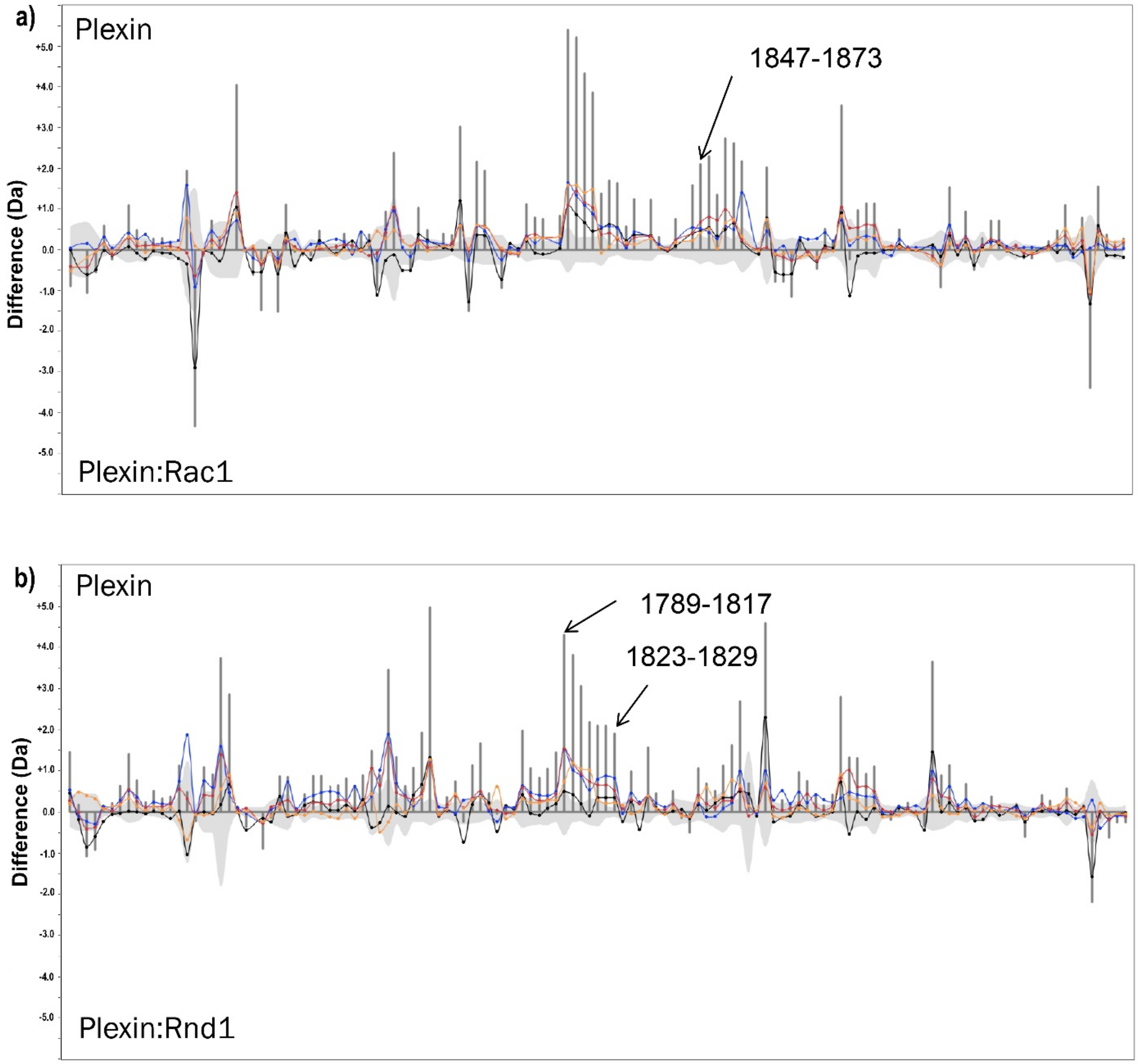

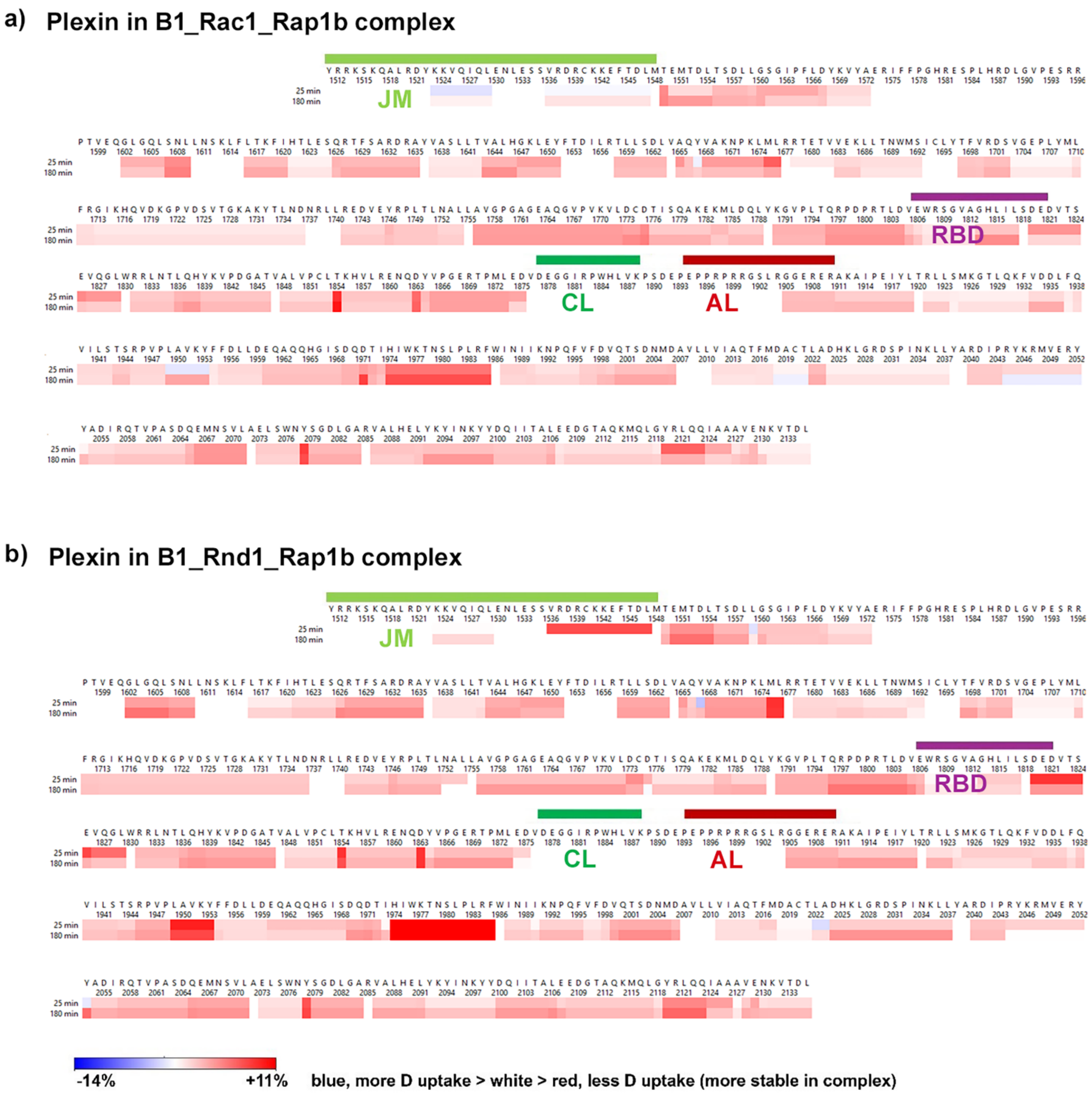


**Supplemental Figure 10: Protection of amides at binding interfaces in plexin when in complex with Rap1b and Rac1 (top) or Rnd1 (bottom):** Protected peptides include most of the JM, RBD-binding and dimerization, and the RBD-GAP interface regions, but only part of the Activation and not the coupling loop. We compared deuterium (D) uptake in plexin-B1 after 25 & 180 min, vs. its uptake in plexin:Rac1:Rap1b and plexin:Rnd1:Rap1b complexes, at pH 7.0, 25 $^{o}$C. The difference data between bound-unbound is shown, plotted on the same scale as in Fig.8. Coupling and Activation loop, as well as RBD position, are indicated above the sequence by green, red and violet, respectively.

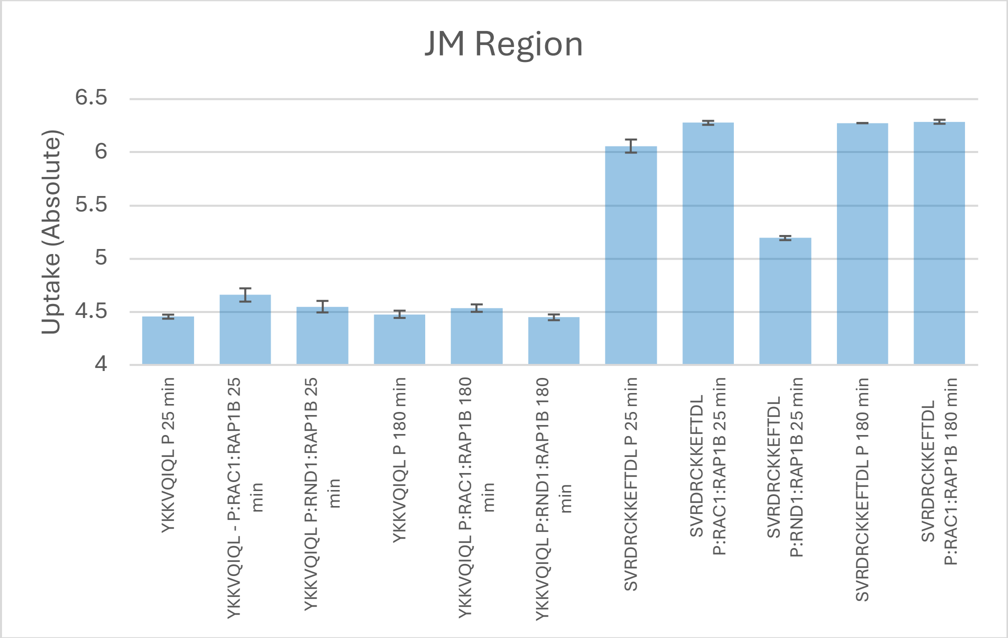


**Supplemental Figure 11. Absolute deuterium uptakes plotted for relevant peptides at 25 or 180 min labelling times within the JM region of interest.** Error bars represent ±1 standard deviation for deuterium uptake.

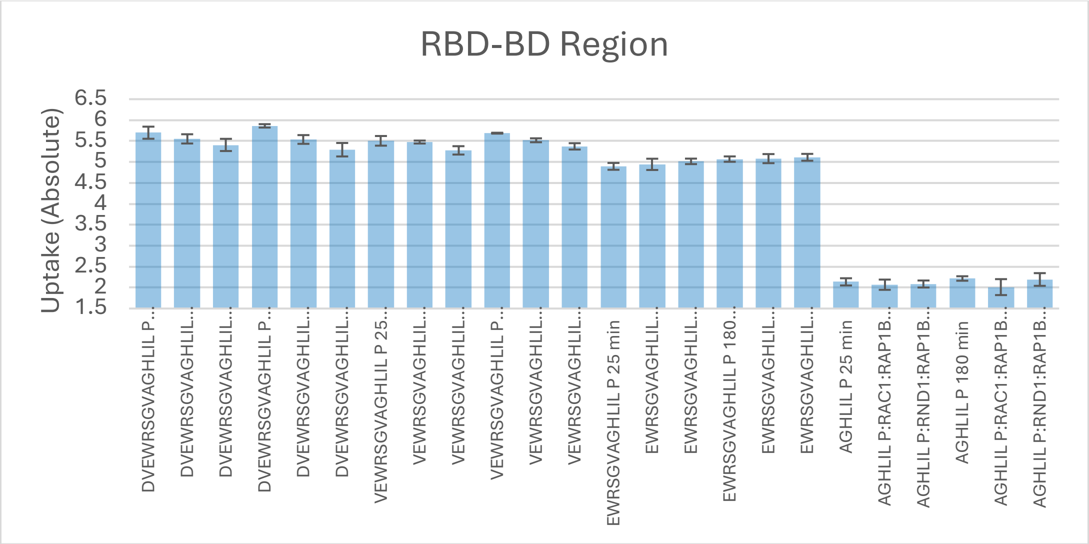

**Supplemental Figure 12. Absolute deuterium uptakes plotted for relevant peptides at 25 or 180 min labelling times within the RBD-BD region of interest.** Error bars represent ±1 standard deviation for deuterium uptake.

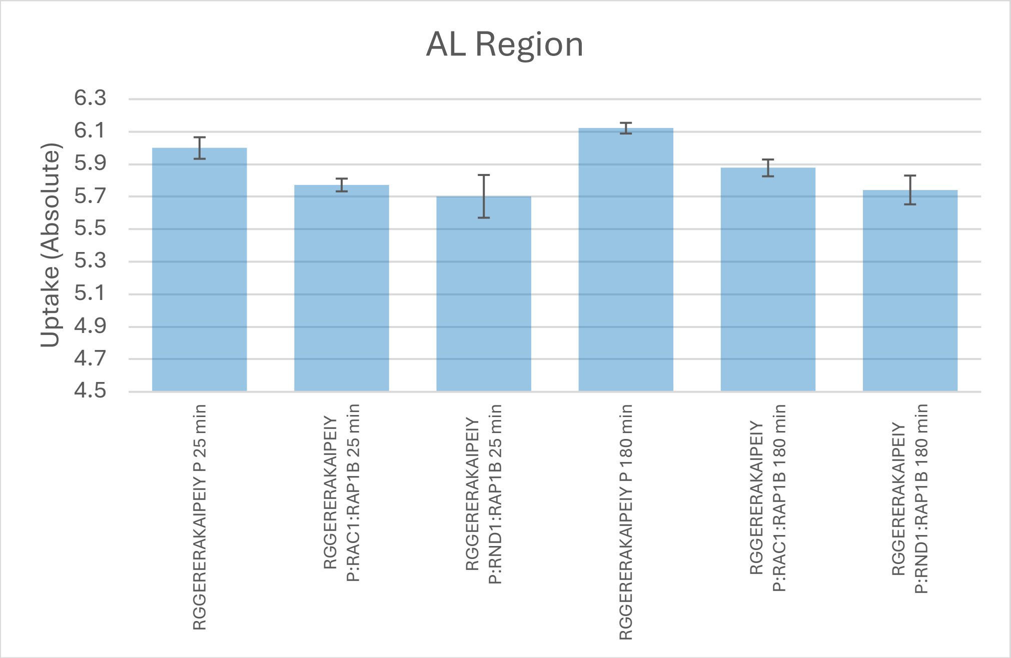


**Supplemental Figure 13. Absolute deuterium uptakes plotted for relevant peptides at 25 or 180 min labelling times within the AL region of interest.** Error bars represent ±1 standard deviation for deuterium uptake.

**Supplemental Table 1. Tabular data for the deuterium uptakes that compose Supplemental Figures 11, 12, and 13 for reference.**

| Region | Start | End | Sequence and State | Exposure (min) | Uptake | Uptake SD | %CV |
|---|---|---|---|---|---|---|---|
| JM | 1522 | 1529 | YKKVQIQL P 25 min | 25 | 4.45 | 0.02 | 0.43 |
| JM | 1522 | 1529 | YKKVQIQL - P:RAC1:RAP1B 25 min | 25 | 4.66 | 0.06 | 1.34 |
| JM | 1522 | 1529 | YKKVQIQL P:RND1:RAP1B 25 min | 25 | 4.55 | 0.05 | 1.19 |
| JM | 1522 | 1529 | YKKVQIQL P 180 min | 180 | 4.48 | 0.03 | 0.76 |
| JM | 1522 | 1529 | YKKVQIQL P:RAC1:RAP1B 180 min | 180 | 4.53 | 0.04 | 0.77 |
| JM | 1522 | 1529 | YKKVQIQL P:RND1:RAP1B 180 min | 180 | 4.45 | 0.03 | 0.61 |
| JM | 1535 | 1547 | SVRDRCKKEFTDL P 25 min | 25 | 6.06 | 0.06 | 1.02 |
| JM | 1535 | 1547 | SVRDRCKKEFTDL P:RAC1:RAP1B 25 min | 25 | 6.28 | 0.02 | 0.30 |
| JM | 1535 | 1547 | SVRDRCKKEFTDL P:RND1:RAP1B 25 min | 25 | 5.19 | 0.02 | 0.38 |
| JM | 1535 | 1547 | SVRDRCKKEFTDL P 180 min | 180 | 6.27 | 0.00 | 0.02 |
| JM | 1535 | 1547 | SVRDRCKKEFTDL P:RAC1:RAP1B 180 min | 180 | 6.29 | 0.02 | 0.30 |
| RBD-BD | 1804 | 1817 | DVEWRSGVAGHLIL P 25 min | 25 | 5.70 | 0.14 | 2.54 |
| RBD-BD | 1804 | 1817 | DVEWRSGVAGHLIL P:RAC1:RAP1B 25 min | 25 | 5.55 | 0.11 | 1.99 |
| RBD-BD | 1804 | 1817 | DVEWRSGVAGHLIL P:RND1:RAP1B 25 min | 25 | 5.41 | 0.15 | 2.69 |

| | | | | | | | |
|---|---|---|---|---|---|---|---|
| RBD-BD | 1804 | 1817 | DVEWRSGVAGHLIL P 180 min | 180 | 5.86 | 0.04 | 0.69 |
| RBD-BD | 1804 | 1817 | DVEWRSGVAGHLIL P:RAC1:RAP1B 180 min | 180 | 5.54 | 0.10 | 1.88 |
| RBD-BD | 1804 | 1817 | DVEWRSGVAGHLIL P:RND1:RAP1B 180 min | 180 | 5.29 | 0.16 | 3.04 |
| RBD-BD | 1805 | 1817 | VEWRSGVAGHLIL P 25 min | 25 | 5.51 | 0.12 | 2.10 |
| RBD-BD | 1805 | 1817 | VEWRSGVAGHLIL P:RAC1:RAP1B 25 min | 25 | 5.48 | 0.04 | 0.64 |
| RBD-BD | 1805 | 1817 | VEWRSGVAGHLIL P:RND1:RAP1B 25 min | 25 | 5.28 | 0.10 | 1.89 |
| RBD-BD | 1805 | 1817 | VEWRSGVAGHLIL P 180 min | 180 | 5.69 | 0.01 | 0.19 |
| RBD-BD | 1805 | 1817 | VEWRSGVAGHLIL P:RAC1:RAP1B 180 min | 180 | 5.52 | 0.05 | 0.85 |
| RBD-BD | 1805 | 1817 | VEWRSGVAGHLIL P:RND1:RAP1B 180 min | 180 | 5.37 | 0.08 | 1.41 |
| RBD-BD | 1806 | 1817 | EWRSGVAGHLIL P 25 min | 25 | 4.90 | 0.08 | 1.67 |
| RBD-BD | 1806 | 1817 | EWRSGVAGHLIL P:RAC1:RAP1B 25 min | 25 | 4.94 | 0.13 | 2.72 |
| RBD-BD | 1806 | 1817 | EWRSGVAGHLIL P:RND1:RAP1B 25 min | 25 | 5.01 | 0.07 | 1.31 |
| RBD-BD | 1806 | 1817 | EWRSGVAGHLIL P 180 min | 180 | 5.07 | 0.06 | 1.26 |
| RBD-BD | 1806 | 1817 | EWRSGVAGHLIL P:RAC1:RAP1B 180 min | 180 | 5.08 | 0.11 | 2.13 |
| RBD-BD | 1806 | 1817 | EWRSGVAGHLIL P:RND1:RAP1B 180 min | 180 | 5.11 | 0.08 | 1.60 |
| RBD-BD | 1812 | 1817 | AGHLIL P 25 min | 25 | 2.14 | 0.09 | 4.03 |
| RBD-BD | 1812 | 1817 | AGHLIL P:RAC1:RAP1B 25 min | 25 | 2.07 | 0.12 | 5.92 |
| RBD-BD | 1812 | 1817 | AGHLIL P:RND1:RAP1B 25 min | 25 | 2.09 | 0.08 | 4.01 |
| RBD-BD | 1812 | 1817 | AGHLIL P 180 min | 180 | 2.22 | 0.05 | 2.42 |
| RBD-BD | 1812 | 1817 | AGHLIL P:RAC1:RAP1B 180 min | 180 | 2.01 | 0.19 | 9.47 |
| RBD-BD | 1812 | 1817 | AGHLIL P:RND1:RAP1B 180 min | 180 | 2.19 | 0.15 | 6.92 |
| AL | 1904 | 1918 | RGGERERAKAIPEIY P 25 min | 25 | 6.00 | 0.07 | 1.10 |
| AL | 1904 | 1918 | RGGERERAKAIPEIY P:RAC1:RAP1B 25 min | 25 | 5.77 | 0.04 | 0.68 |
| AL | 1904 | 1918 | RGGERERAKAIPEIY P:RND1:RAP1B 25 min | 25 | 5.70 | 0.13 | 2.31 |
| AL | 1904 | 1918 | RGGERERAKAIPEIY P 180 min | 180 | 6.12 | 0.03 | 0.54 |
| AL | 1904 | 1918 | RGGERERAKAIPEIY P:RAC1:RAP1B 180 min | 180 | 5.88 | 0.05 | 0.88 |
| AL | 1904 | 1918 | RGGERERAKAIPEIY P:RND1:RAP1B 180 min | 180 | 5.74 | 0.09 | 1.54 |
